# CB_1_ Cannabinoid Receptor Agonists Induce Acute Respiratory Depression in Awake Mice

**DOI:** 10.1101/2024.03.12.584260

**Authors:** Joshua Watkins, Petra Aradi, Rachel Hahn, Istvan Katona, Ken Mackie, Alexandros Makriyannis, Andrea G. Hohmann

## Abstract

Recreational use of synthetic cannabinoid agonists (i.e., “Spice” compounds) that target the Cannabinoid Type 1 receptor (CB_1_) can cause respiratory depression in humans. However, Δ^9^-tetrahydrocannabinol (THC), the major psychoactive phytocannabinoid in cannabis, is not traditionally thought to interact with CNS control of respiration, based largely upon sparse labeling of CB1 receptors in the medulla and few reports of clinically significant respiratory depression following cannabis overdose. The respiratory effects of CB_1_ agonists have rarely been studied *in vivo*, suggesting that additional inquiry is required to reconcile the conflict between conventional wisdom and human data. Here we used whole body plethysmography to examine the respiratory effects of the synthetic high efficacy CB_1_ agonist CP55,940, and the low efficacy CB_1_ agonist Δ^9^-tetrahydrocannabinol in male and female mice. CP55,940 and THC, administered systemically, both robustly suppressed minute ventilation. Both cannabinoids also produced sizable reductions in tidal volume, decreasing both peak inspiratory and expiratory flow - measures of respiratory effort. Similarly, both drugs reduced respiratory frequency, decreasing both inspiratory and expiratory time while markedly increasing expiratory pause, and to a lesser extent, inspiratory pause. Respiratory suppressive effects occurred at lower doses in females than in males, and at many of the same doses shown to produce cardinal behavioral signs of CB_1_ activation. We next used RNAscope *in situ* hybridization to localize CB_1_ mRNA to glutamatergic neurons in the medullary pre-Bötzinger Complex, a critical nucleus in controlling respiration. Our results show that, contrary to previous conventional wisdom, CB_1_ mRNA is expressed in glutamatergic neurons in a brain region essential for breathing and CB_1_ agonists can cause significant respiratory depression.

## 1. Introduction

Despite widespread human use of cannabis, reports of lethal overdose are rare. Respiratory depression, the predominant cause of death in drug overdose, is largely absent in users of cannabis; there are no reports of human cannabis use leading to death by respiratory depression. Respiratory depression after exogenous drug administration (e.g., opioids) results from inhibition of the neural systems that control breathing [1, 2]. Because the main psychoactive component of cannabis, Δ^9^-tetrahydrocannabinol (THC), acts as a weak partial agonist of the Cannabinoid Type 1 (CB_1_) receptor [3–9], this lack of lethal overdoses has led to the perception that cannabinoid drugs, in general, do not strongly interact with the respiratory system. This position is bolstered by receptor binding and autoradiography studies demonstrating that expression of cannabinoid receptors is sparse in the medulla, where breathing is primarily controlled [10].

This perception of relative safety has been challenged by the emergence of illicit synthetic drugs targeting the CB_1_ receptor. These compounds, sold under generic names (e.g., “Spice”, “K2”, synthetic marijuana, etc.) and marketed as legal or more potent alternatives to cannabis, are typically high-potency and high-efficacy agonists of the CB_1_ receptor [11–14]. In stark contrast to phytocannabinoid products, clinical literature surrounding the use of these illicit synthetic cannabinoids is replete with case reports in which users suffer acute respiratory depression requiring emergency medical care, including intubation and mechanical ventilation [15–21]. Furthermore, case reports have identified synthetic cannabinoid use in patients with opioid use disorder, and coadministration of synthetic cannabinoids and synthetic opioids in lethal overdose [22, 23]. This emerging clinical literature requires that the possible role and distribution of cannabinoid receptors within respiratory circuits be reevaluated using the most sensitive techniques available.

Here we present evidence that the prototypical cannabinoid agonists CP55,940 and THC induce respiratory depression in awake adult mice of both sexes after IP injection. We evaluated the ability of the synthetic CB_1_ cannabinoid full agonist CP55,940 and CB_1_ partial agonist Δ^9^-tetrahydrocannabinol for their ability to produce respiratory depression in wild type mice. Pharmacological specificity was assessed using both brain penetrant (AM251) and peripherally restricted (AM6545) CB_1_ antagonists. Finally, we assessed the distribution of CB_1_ receptor mRNA in brainstem regions that are known to control respiratory activity using RNAscope *in situ* hybridization and identified phenotypes of CB_1_ mRNA-expressing cells, demonstrating that CB_1_ mRNA is expressed in the pre-Bötzinger Complex (preBötC), a brain region known to be critical to respiration [24–27].

## 2. Materials and Methods

### 2.1. Subjects

Subjects consisted of 125 male and 70 female C57BL/6J wild type mice (Jackson Laboratories; Bar Harbor, ME; 12 weeks of age). Male mice weighed 30-40g, and female mice weighed 19-30g prior to testing. Mice were housed in a temperature- and humidity-controlled facility with *ad libitum* access to food and water on a 12:12h light/dark cycle. All procedures were approved by the Bloomington Institutional Animal Care and Use Committee of Indiana University.

### 2.2. Drugs

CP55,940 was purchased from Cayman Pharmaceuticals (Ann Arbor, MI). THC was obtained from the NIDA Drug Supply Program (Division of Therapeutics and Medical Consequences, National Institutes of Health). AM251 and AM6545 were synthesized by the Center for Drug Discovery (Northeastern University, Boston, MA).

All drugs used in pharmacological experiments were administered intraperitoneally in a volume of 10 ml/kg. Drugs were administered in a vehicle consisting of 10% dimethylsulfoxide (DMSO; Sigma-Aldrich, St. Louis, MO) with the remaining volume consisting of emulphor (Alkamuls EL-620; Solvay), 100% ethanol (Decon Labs; Montgomery, PA), and 0.9% saline (Baxter Pharmaceuticals; Bloomington, IN) at a 1:1:18 ratio except where noted. Due to limited solubility, in experiments involving AM6545, all drugs were administered in a vehicle consisting of 20% DMSO, with the remaining volume consisting of emulphor, ethanol, and saline at a 1:1:18 ratio.

### 2.3. Assessment of respiratory parameters

Whole body plethysmography (Data Sciences International; St. Paul, MN) was used to assess respiratory parameters in awake unrestrained mice. Evaluated parameters were respiratory frequency (breaths per minute), tidal volume (volume of individual breaths), minute ventilation (volume of ventilation in mL/minute), peak inspiratory flow (mL/second), peak expiratory flow (mL/second), end inspiratory pause (time between inspiration and expiration; ms), and end expiratory pause (time between inspiratory cycles; ms). Animals were habituated to recording chambers for 30 minutes within 48 hours of testing to minimize changes in respiration resulting from stress related to a novel environment. On test day mice received air mixed with 10% CO_2_ to eliminate hypercapnic responses without inducing stress. Testing consisted of a 50-minute initial recording to acquire baseline respiratory parameters, after which mice were briefly removed from test chambers and given a single IP injection containing drug. Animals were then returned to the chambers for 60 minutes to obtain post-dose respiratory parameters.

### 2.4. Perfusion

Naive male CD1 wild type and CB_1_^−/−^ mice (postnatal day 84; n = 5 per group) were fixed via transcardial perfusion. Mice were deeply anesthetized with isoflurane (Covetrus, 11695-6777-2) and perfused transcardially with 0.9% saline for 2 min, following with 4% paraformaldehyde (PFA, Sigma, P6148) dissolved in 0.1 M phosphate buffer (PB, pH 7.4 containing Na2HPO4, Sigma, 71 643 and NaH2PO4, Sigma, 71 500) for 18 min by using approximately 5 mL/min pump speed. After perfusion, the brains were removed from the skull and postfixed at room temperature overnight in 4% PFA, then 30 μm coronal sections were cut in PB using a Leica VT-1200S vibratome (Nussloch, Germany). All solutions were prepared from filtered water distilled through a Biopak Polisher (Sigma, CDUFBI001) to decrease RNAse contamination.

### 2.5. RNAscope *in situ* hybridization

RNAscope *in situ* hybridization was used to detect cell type-specific CB_1_ mRNA expression in CB_1_^−/−^ mice and their wild type littermates. Free-floating tissue sections were mounted on positively charged microscopic Superfrost Plus slides (Fisher Scientific), then a series of dehydration steps were performed using 50%, 70%, and 100% ethanol (Sigma, E7023), and sections were incubated in 100% ethanol overnight at 4C. The next day, the sections were dried, and a hydrophobic barrier was drawn around the slices with a hydrophobic pen (ImmEdge Hydrophobic Barrier, Vector Laboratories). RNAscope mRNA-staining steps were performed following the manufacturer’s protocols. Stained slides were coverslipped with fluorescent mounting medium (ProLong Diamond Anti-fade Reagent P36961; Invitrogen) and imaged with Nikon A1 confocal laser-scanning microscope. Analysis of *in situ* hybridization was completed using QuPath (v.0.5.0) subcellular detection.

Gene-specific RNAscope probes were used to target the following mRNA sequences: CB_1_ (catalog #530-1458, targeting 530-1458 bp of the mouse Cnr1 sequence, NM_007726.3), VGLUT2 (catalog #319171-C3, targeting 1986-2998 bp of the mouse VGLUT2 (Slc17a6) sequence, NM_080853.3) and ChAT (catalog #408731-C2, targeting 1090-1952 bp of the mouse Chat sequence, NM_009891.2). All probes were designed and provided by Advanced Cell Diagnostics (Newark, CA).

### 2.6. Statistical analysis

All statistics were calculated and tested using GraphPad Prism version 9.4.0. Data was averaged into five-minute bins and represented as mean ± SEM. Two-way repeated measures ANOVA was used to assess respiratory effects of all pharmacological manipulations across time. Dunnett’s post hoc test was used to compare main effects of drug treatment with vehicle controls. Bonferroni’s multiple comparison post hoc test was used to assess the impact of antagonist treatments on respiratory parameters. Pre-injection and post-injection respiratory parameters were measured separately to ensure that differences in baseline responsiveness did not impact interpretation of drug effects. Dose response curves were generated by logarithmically transforming the difference between the final pre- and post-dose bin and fitting a four-parameter logistic regression model.

## 3. Results

### General results

Prior to pharmacological manipulations, respiratory parameters declined across time, but at the timepoint of drug injection there were no group differences as a function of drug allocation except where noted. Observed effects were rapid and persistent across the entire recording session; once depressed, no metrics recovered to control levels for the remaining duration of recording. Despite the magnitude and persistence of observed respiratory depression, no fatalities were observed in experimental animals.

### 3.1. Respiratory effects of CP55,940

CP55,940 markedly decreased minute ventilation in a dose- and time-dependent manner. Post-hoc comparisons revealed that all doses 0.1 mg/kg and above suppressed minute ventilation in both males and females [Figure 1; Tables 1 & 2]. This suppression of overall respiratory activity was driven by a reduction to both the number of initiated breaths and the amplitude of gas exchange of each breath; CP55,940 suppressed both respiratory frequency [Figure 2] and tidal volume [Figure 3] in a dose- and time-dependent manner. Frequency and tidal volume were both suppressed by all doses 0.1 mg/kg and above in animals of both sexes.

**Figure 1:**
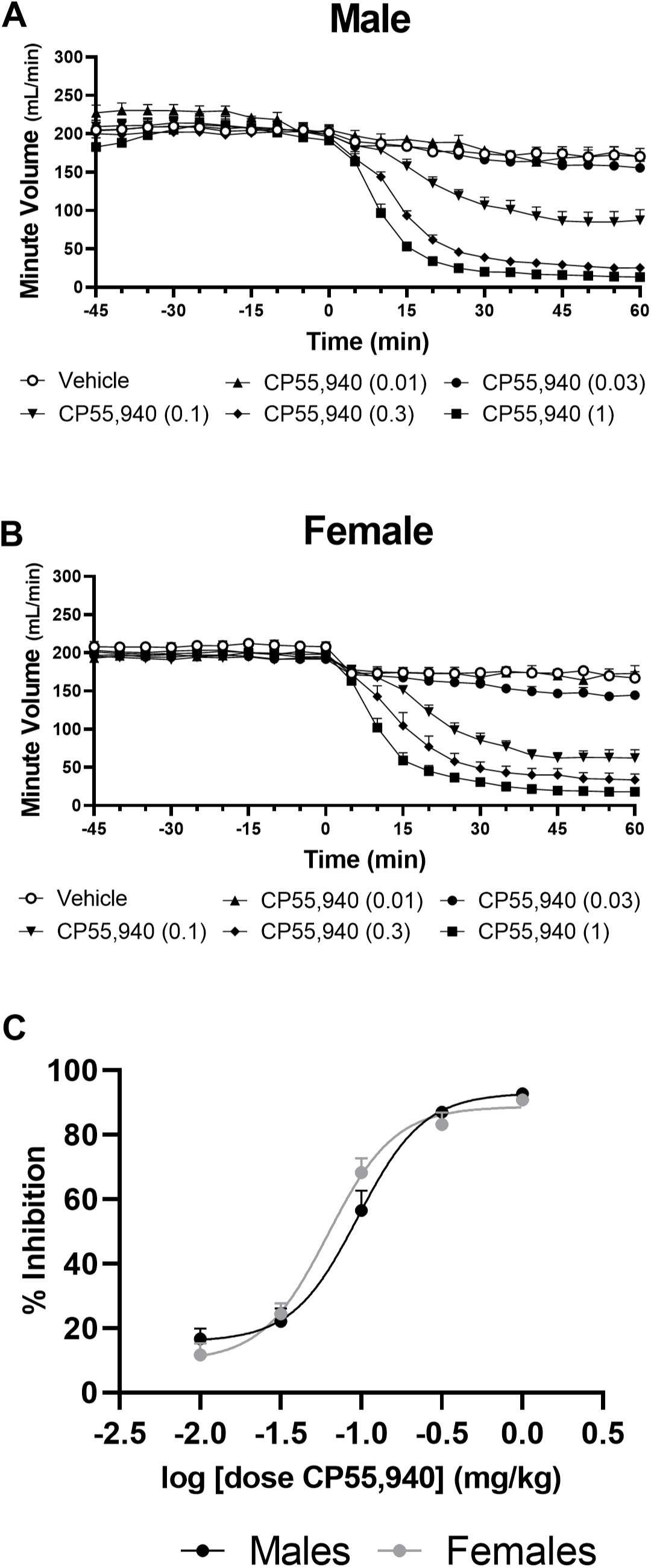
CP55,940 Minute Ventilation and Dose Response

**Figure 2:**
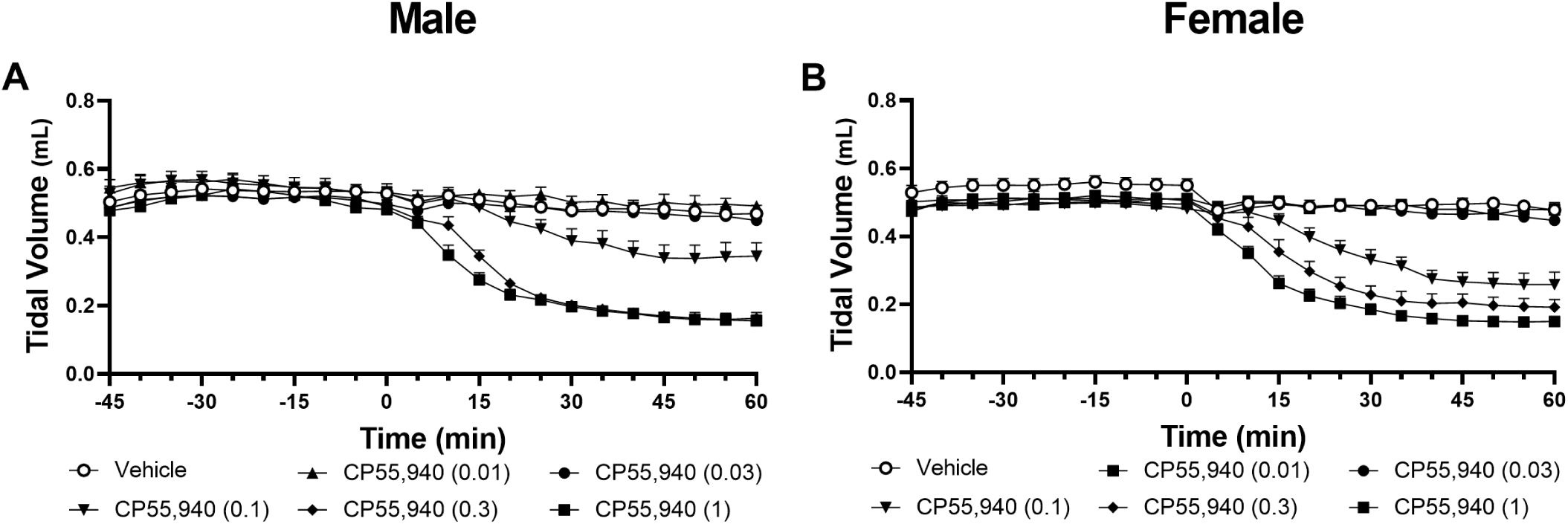
CP55,940 Tidal Volume

**Figure 3:**
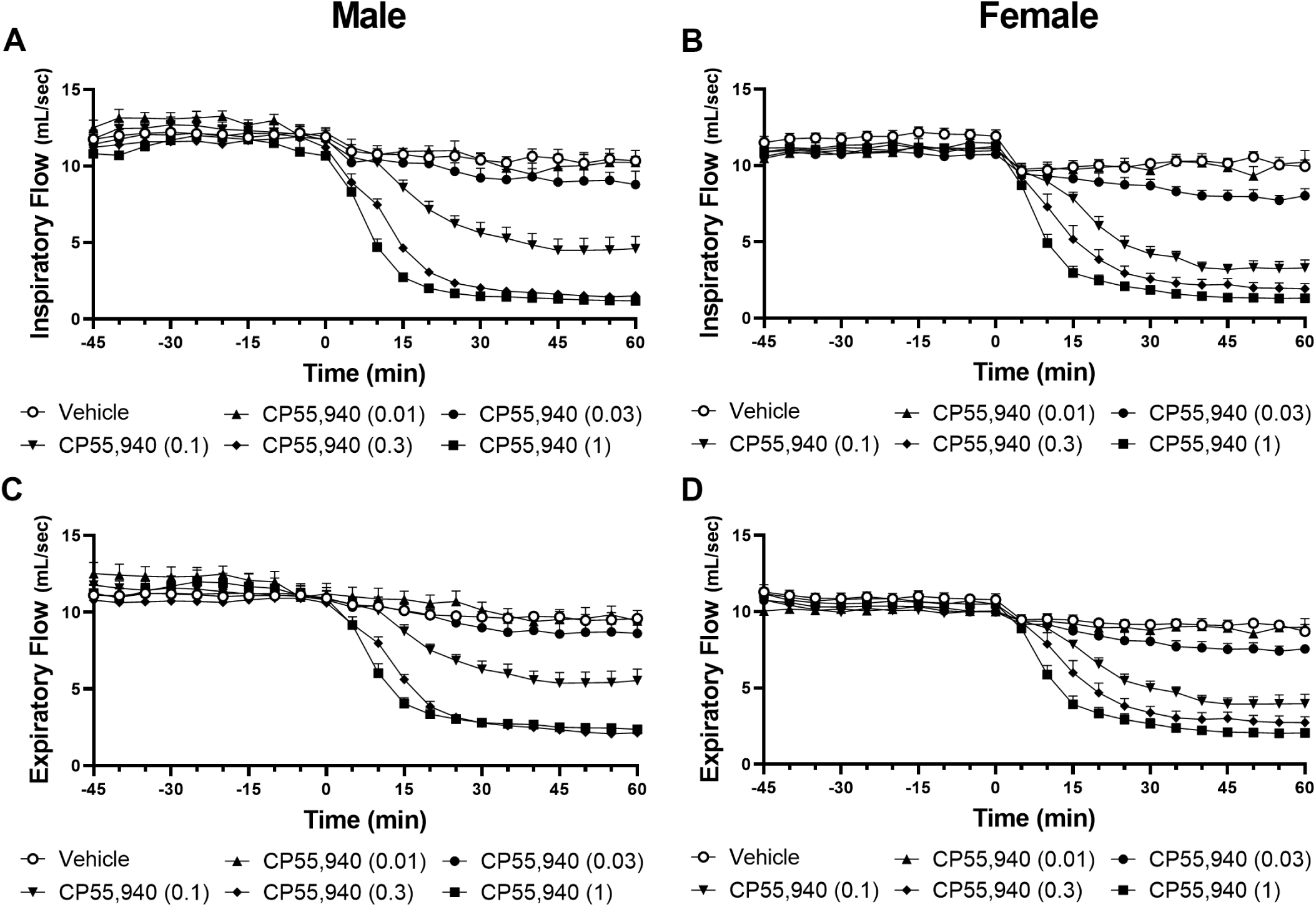
CP55,940 Inspiratory and Expiratory Flow

**Table 1:** CP55,940 Males.

| Metric | Time |  | Drug |  | Interaction |  | Effective Doses (mg/kg) |  |  |  |  |
| --- | --- | --- | --- | --- | --- | --- | --- | --- | --- | --- | --- |
|  |  | p |  | p |  | p | 0.01 | 0.03 | 0.1 | 0.3 | 1 |
| Minute Ventilation | $F_{11,363} = 120.1$ | 0.0001 | $F_{5,33} = 120.6$ | 0.0001 | $F_{55,363} = 34.85$ | 0.0001 | - | - | 0.0001 | 0.0001 | 0.0001 |
| Tidal Volume | $F_{11,363} = 58.91$ | 0.0001 | $F_{5,33} = 138.9$ | 0.0001 | $F_{55,363} = 23.70$ | 0.0001 | - | - | 0.0200 | 0.0001 | 0.0001 |
| Inspiratory Flow | $F_{11,363} = 130.7$ | 0.0001 | $F_{5,33} = 74.7$ | 0.0001 | $F_{55,363} = 16.46$ | 0.0001 | - | - | 0.0001 | 0.0001 | 0.0001 |
| Expiratory Flow | $F_{11,363} = 192.0$ | 0.0001 | $F_{5,33} = 52.94$ | 0.0001 | $F_{55,363} = 17.85$ | 0.0001 | - | - | 0.0001 | 0.0001 | 0.0001 |
| Frequency | $F_{11,363} = 249.5$ | 0.0001 | $F_{5,33} = 163.5$ | 0.0001 | $F_{55,363} = 38.21$ | 0.0001 | - | - | 0.0001 | 0.0001 | 0.0001 |
| Inspiratory Time | $F_{11,363} = 120.1$ | 0.0001 | $F_{5,33} = 120.6$ | 0.0001 | $F_{55,363} = 34.85$ | 0.0001 | - | - | 0.0015 | 0.0001 | 0.0001 |
| Expiratory Time | $F_{11,363} = 58.91$ | 0.0001 | $F_{5,33} = 138.9$ | 0.0001 | $F_{55,363} = 23.70$ | 0.0001 | - | - | - | 0.0001 | 0.0001 |
| Inspiratory Pause | $F_{11,363} = 91.73$ | 0.0001 | $F_{5,33} = 185.9$ | 0.0001 | $F_{55,363} = 46.67$ | 0.0001 | - | - | - | 0.0001 | 0.0001 |
| Expiratory Pause | $F_{11,363} = 15.37$ | 0.0001 | $F_{5,33} = 29.58$ | 0.0001 | $F_{55,363} = 13.00$ | 0.0001 | - | - | - | - | 0.0001 |

**Table 2:**
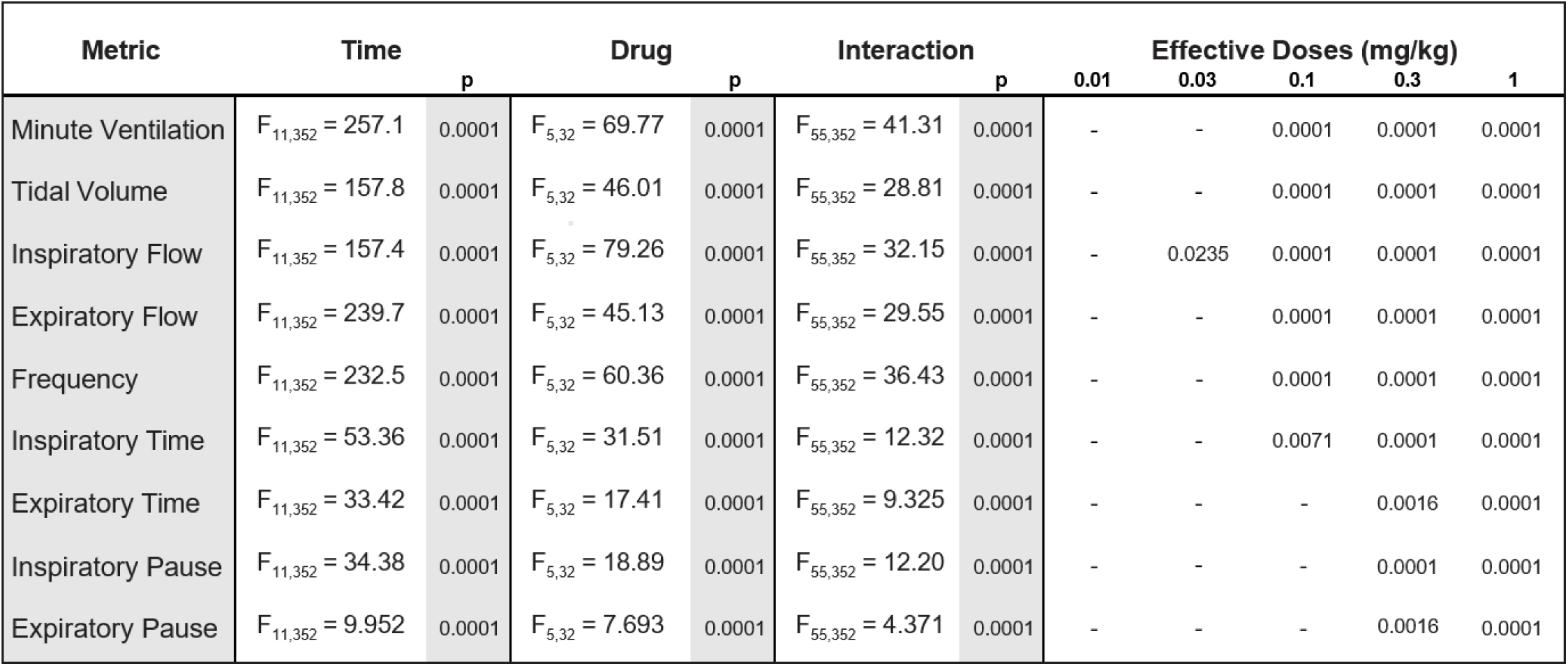
CP55,940 Females.

| Metric | Time |  | Drug |  | Interaction |  | Effective Doses (mg/kg) |  |  |  |  |
| --- | --- | --- | --- | --- | --- | --- | --- | --- | --- | --- | --- |
|  |  | p |  | p |  | p | 0.01 | 0.03 | 0.1 | 0.3 | 1 |
| Minute Ventilation | $F_{11,352} = 257.1$ | 0.0001 | $F_{5,32} = 69.77$ | 0.0001 | $F_{55,352} = 41.31$ | 0.0001 | - | - | 0.0001 | 0.0001 | 0.0001 |
| Tidal Volume | $F_{11,352} = 157.8$ | 0.0001 | $F_{5,32} = 46.01$ | 0.0001 | $F_{55,352} = 28.81$ | 0.0001 | - | - | 0.0001 | 0.0001 | 0.0001 |
| Inspiratory Flow | $F_{11,352} = 157.4$ | 0.0001 | $F_{5,32} = 79.26$ | 0.0001 | $F_{55,352} = 32.15$ | 0.0001 | - | 0.0235 | 0.0001 | 0.0001 | 0.0001 |
| Expiratory Flow | $F_{11,352} = 239.7$ | 0.0001 | $F_{5,32} = 45.13$ | 0.0001 | $F_{55,352} = 29.55$ | 0.0001 | - | - | 0.0001 | 0.0001 | 0.0001 |
| Frequency | $F_{11,352} = 232.5$ | 0.0001 | $F_{5,32} = 60.36$ | 0.0001 | $F_{55,352} = 36.43$ | 0.0001 | - | - | 0.0001 | 0.0001 | 0.0001 |
| Inspiratory Time | $F_{11,352} = 53.36$ | 0.0001 | $F_{5,32} = 31.51$ | 0.0001 | $F_{55,352} = 12.32$ | 0.0001 | - | - | 0.0071 | 0.0001 | 0.0001 |
| Expiratory Time | $F_{11,352} = 33.42$ | 0.0001 | $F_{5,32} = 17.41$ | 0.0001 | $F_{55,352} = 9.325$ | 0.0001 | - | - | - | 0.0016 | 0.0001 |
| Inspiratory Pause | $F_{11,352} = 34.38$ | 0.0001 | $F_{5,32} = 18.89$ | 0.0001 | $F_{55,352} = 12.20$ | 0.0001 | - | - | - | 0.0001 | 0.0001 |
| Expiratory Pause | $F_{11,352} = 9.952$ | 0.0001 | $F_{5,32} = 7.693$ | 0.0001 | $F_{55,352} = 4.371$ | 0.0001 | - | - | - | 0.0016 | 0.0001 |

Observed changes to frequency were driven by an extension of all respiratory phases, though the doses at which these effects were observed were not uniform. Doses of CP55,940 0.1 mg/kg and above extended inspiratory time in animals of both sexes, while expiratory time and inspiratory pause were extended by doses of 0.3 mg/kg and above in both sexes [Figure 4]. End expiratory pause was extended by doses 0.3 mg/kg and above in females, and only by the highest tested dose, 1 mg/kg, in males. Changes to tidal volume were driven by both inspiratory and expiratory effort. Inspiratory flow was suppressed by all doses 0.1 mg/kg and higher in males, and all doses 0.03 mg/kg and higher in females, while expiratory flow was suppressed by all doses 0.1 mg/kg and higher in both sexes [Figure 5].

**Figure 4:**
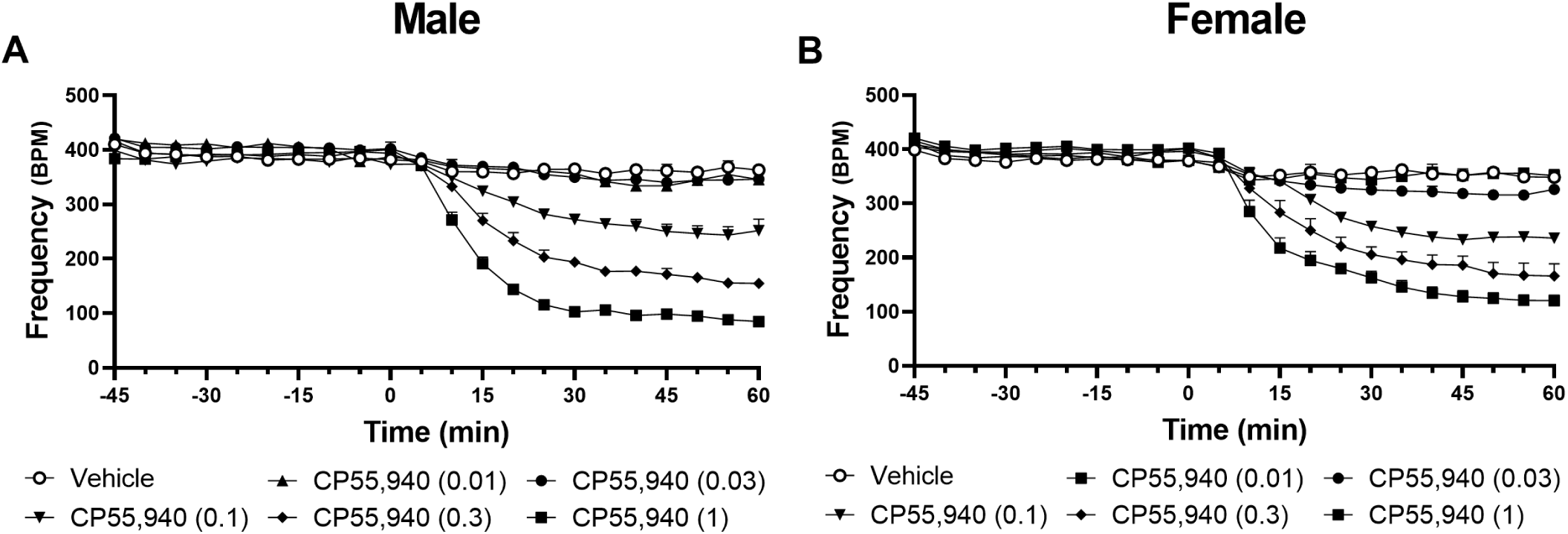
CP55,940 Frequency

**Figure 5:**
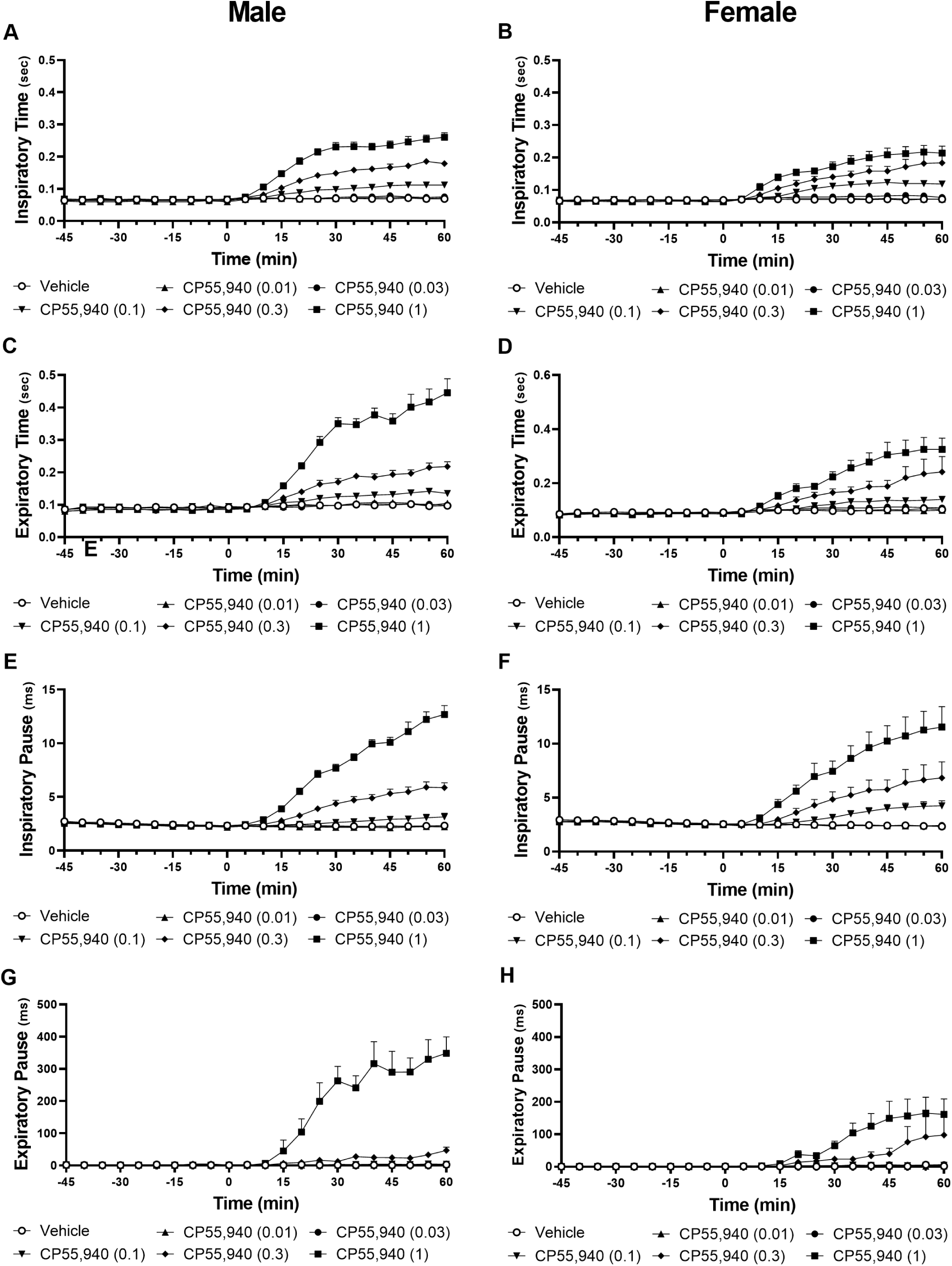
CP55,940 Inspiratory Time, Expiratory Time, End Inspiratory Pause, and End Expiratory Pause

### 3.2. Respiratory effects of THC

THC also markedly decreased minute ventilation in a dose- and time-dependent manner, though these effects occurred at higher doses than CP55,940. Doses 3 mg/kg and above suppressed minute ventilation in males, and 1 mg/kg suppressed minute ventilation in females [Figure 6; Tables 3 & 4]. As with CP55,940, this effect was driven by reductions to both respiratory frequency and tidal volume. In males, all doses 1 mg/kg and above suppressed frequency [Figure 7], while 10 mg/kg and 30 mg/kg doses suppressed tidal volume [Figure 8]. In females, all doses 3 mg/kg and above suppressed both frequency and tidal volume.

**Figure 6:**
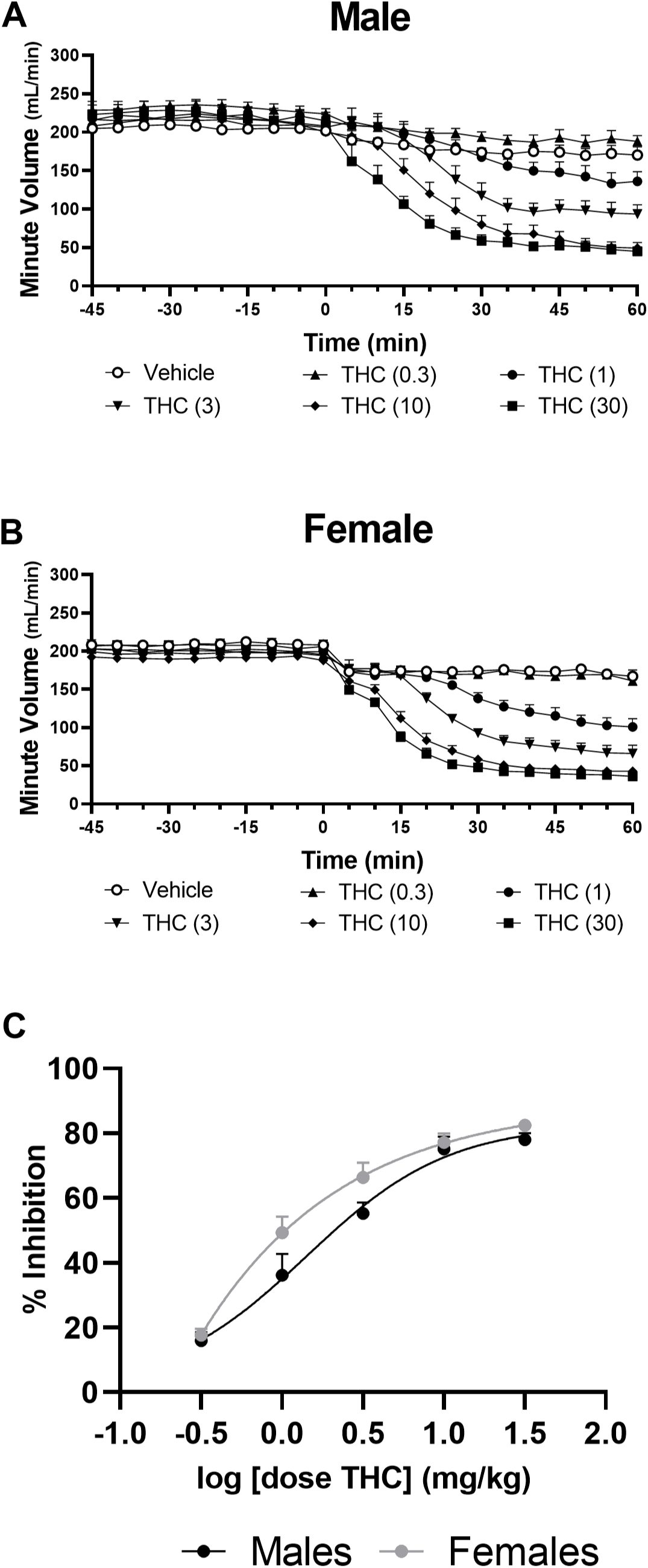
THC Minute Ventilation and Dose Response

**Figure 7:**
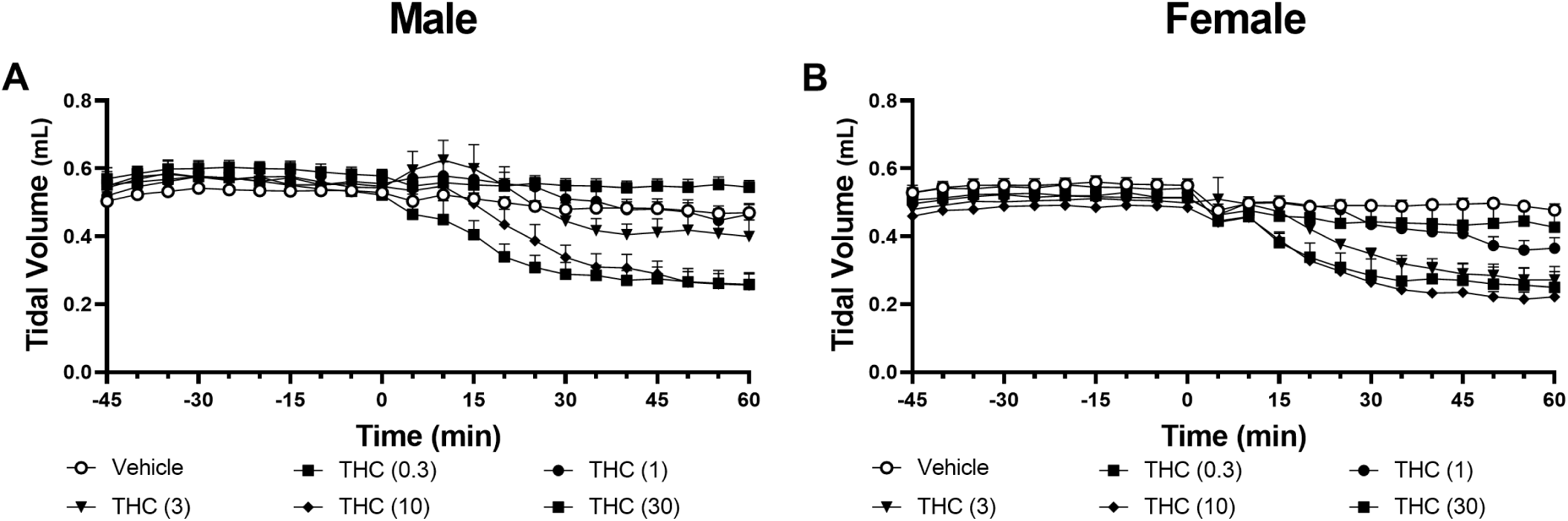
THC Tidal Volume

**Figure 8:**
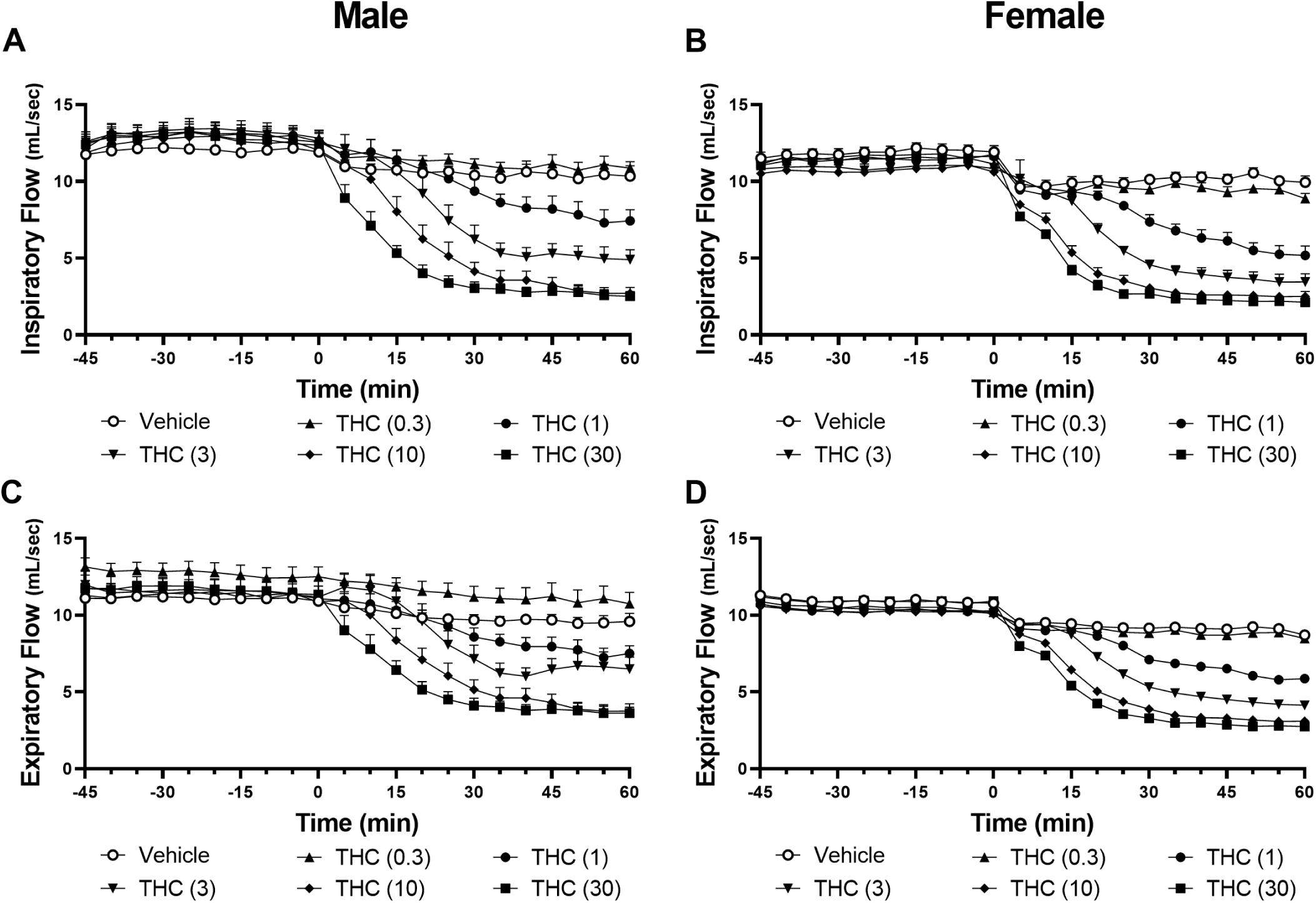
THC Inspiratory and Expiratory Flow

**Table 3:** THC Males.

| Metric | Time |  | Drug |  | Interaction |  | Effective Doses (mg/kg) |  |  |  |  |
| --- | --- | --- | --- | --- | --- | --- | --- | --- | --- | --- | --- |
|  |  | p |  | p |  | p | 0.3 | 1 | 3 | 10 | 30 |
| Minute Ventilation | $F_{11,352} = 169.5$ | 0.0001 | $F_{5,32} = 23.80$ | 0.0001 | $F_{55,352} = 13.81$ | 0.0001 | - | - | 0.0146 | 0.0001 | 0.0001 |
| Tidal Volume | $F_{11,352} = 103.9$ | 0.0049 | $F_{5,32} = 7.667$ | 0.0001 | $F_{55,352} = 10.56$ | 0.0001 | - | - | - | 0.0403 | 0.0022 |
| Inspiratory Flow | $F_{11,352} = 142.6$ | 0.0001 | $F_{5,32} = 27.95$ | 0.0001 | $F_{55,352} = 14.49$ | 0.0001 | - | - | 0.0008 | 0.0001 | 0.0001 |
| Expiratory Flow | $F_{11,352} = 183.5$ | 0.0001 | $F_{5,32} = 17.14$ | 0.0001 | $F_{55,352} = 13.93$ | 0.0001 | - | - | - | 0.0001 | 0.0001 |
| Frequency | $F_{11,352} = 240.0$ | 0.0001 | $F_{5,32} = 44.90$ | 0.0001 | $F_{55,352} = 21.85$ | 0.0001 | - | 0.0217 | 0.0001 | 0.0001 | 0.0001 |
| Inspiratory Time | $F_{11,352} = 147.7$ | 0.0001 | $F_{5,32} = 49.97$ | 0.0001 | $F_{55,352} = 23.29$ | 0.0001 | - | - | 0.0001 | 0.0001 | 0.0001 |
| Expiratory Time | $F_{11,352} = 103.1$ | 0.0001 | $F_{5,32} = 22.72$ | 0.0001 | $F_{55,352} = 13.13$ | 0.0001 | - | - | 0.0015 | 0.0001 | 0.0001 |
| Inspiratory Pause | $F_{11,352} = 76.33$ | 0.0001 | $F_{5,32} = 21.84$ | 0.0001 | $F_{55,352} = 20.95$ | 0.0001 | - | - | - | 0.0001 | 0.0001 |
| Expiratory Pause | $F_{11,352} = 9.267$ | 0.0001 | $F_{5,32} = 6.863$ | 0.0002 | $F_{55,352} = 2.875$ | 0.0002 | - | - | - | 0.0019 | 0.0036 |

**Table 4:** THC Females.

| Metric | Time |  | Drug |  | Interaction |  | Effective Doses (mg/kg) |  |  |  |  |
| --- | --- | --- | --- | --- | --- | --- | --- | --- | --- | --- | --- |
|  |  | p |  | p |  | p | 0.3 | 1 | 3 | 10 | 30 |
| Minute Ventilation | $F_{11,352} = 239.6$ | 0.0001 | $F_{5,32} = 81.49$ | 0.0001 | $F_{55,352} = 28.71$ | 0.0001 | - | 0.0001 | 0.0001 | 0.0001 | 0.0001 |
| Tidal Volume | $F_{11,352} = 80.57$ | 0.0001 | $F_{5,32} = 9.128$ | 0.0001 | $F_{55,352} = 8.919$ | 0.0001 | - | - | 0.0063 | 0.0001 | 0.0001 |
| Inspiratory Flow | $F_{11,352} = 152.6$ | 0.0001 | $F_{5,32} = 92.12$ | 0.0001 | $F_{55,352} = 22.75$ | 0.0001 | - | 0.0001 | 0.0001 | 0.0001 | 0.0001 |
| Expiratory Flow | $F_{11,352} = 182.0$ | 0.0001 | $F_{5,32} = 52.68$ | 0.0001 | $F_{55,352} = 16.95$ | 0.0001 | - | 0.0005 | 0.0001 | 0.0001 | 0.0001 |
| Frequency | $F_{11,352} = 152.5$ | 0.0001 | $F_{5,32} = 14.13$ | 0.0001 | $F_{55,352} = 12.82$ | 0.0001 | - | - | 0.0045 | 0.0001 | 0.0001 |
| Inspiratory Time | $F_{11,352} = 158.0$ | 0.0001 | $F_{5,32} = 92.31$ | 0.0001 | $F_{55,352} = 23.18$ | 0.0001 | - | 0.0009 | 0.0001 | 0.0001 | 0.0001 |
| Expiratory Time | $F_{11,352} = 148.7$ | 0.0001 | $F_{5,32} = 41.34$ | 0.0001 | $F_{55,352} = 20.03$ | 0.0001 | - | - | 0.0116 | 0.0001 | 0.0001 |
| Inspiratory Pause | $F_{11,352} = 92.81$ | 0.0001 | $F_{5,32} = 45.40$ | 0.0001 | $F_{55,352} = 22.17$ | 0.0001 | - | - | 0.0010 | 0.0001 | 0.0001 |
| Expiratory Pause | $F_{11,352} = 16.07$ | 0.0001 | $F_{5,32} = 8.021$ | 0.0001 | $F_{55,352} = 3.897$ | 0.0001 | - | - | - | - | 0.0001 |

Inspiratory time was extended by all doses 3 mg/kg and above in males and 1 mg/kg and above in females, while expiratory time was increased by all doses 3 mg/kg and above in both sexes [Figure 9]. Expiratory pause was extended by doses 10 mg/kg and above in males, and only at the highest tested dose of 30 mg/kg in females. Inspiratory pause was extended by all doses 3 mg/kg and above in females, and by all doses 10 mg/kg and above in males. Inspiratory flow was suppressed by doses 3 mg/kg and above in males, and 1 mg/kg and above in females, while expiratory flow was suppressed by doses 10 mg/kg and above in males and 1 mg/kg and above in females [Figure 10].

**Figure 9:**
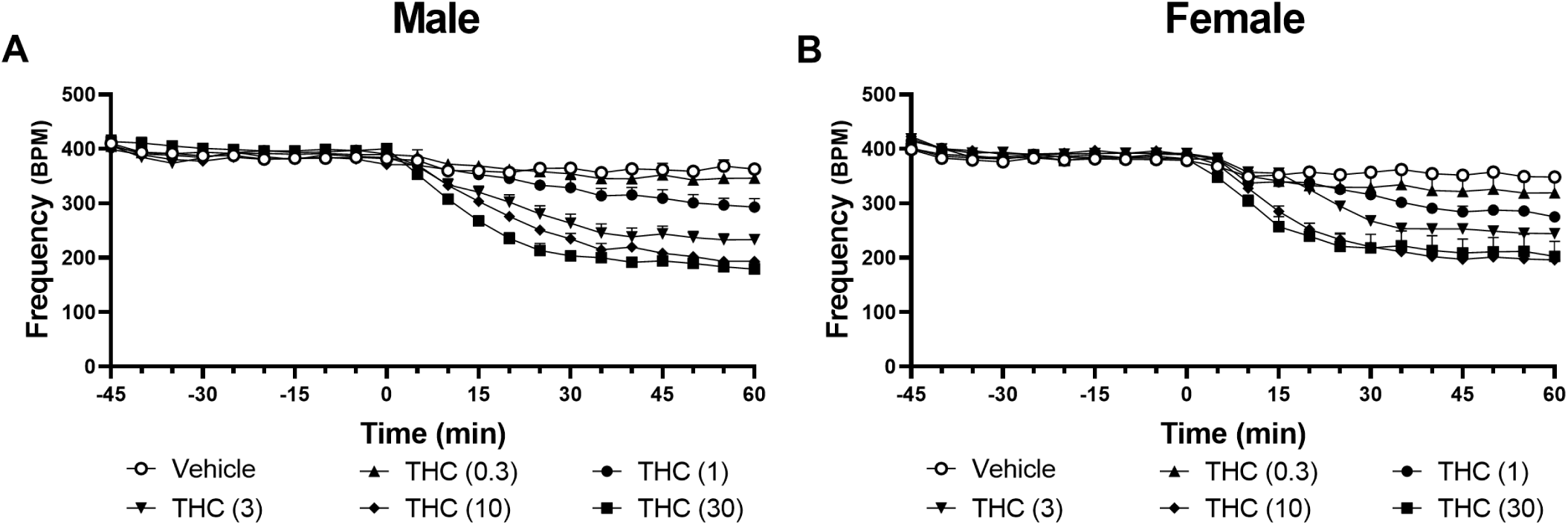
THC Frequency

**Figure 10:**
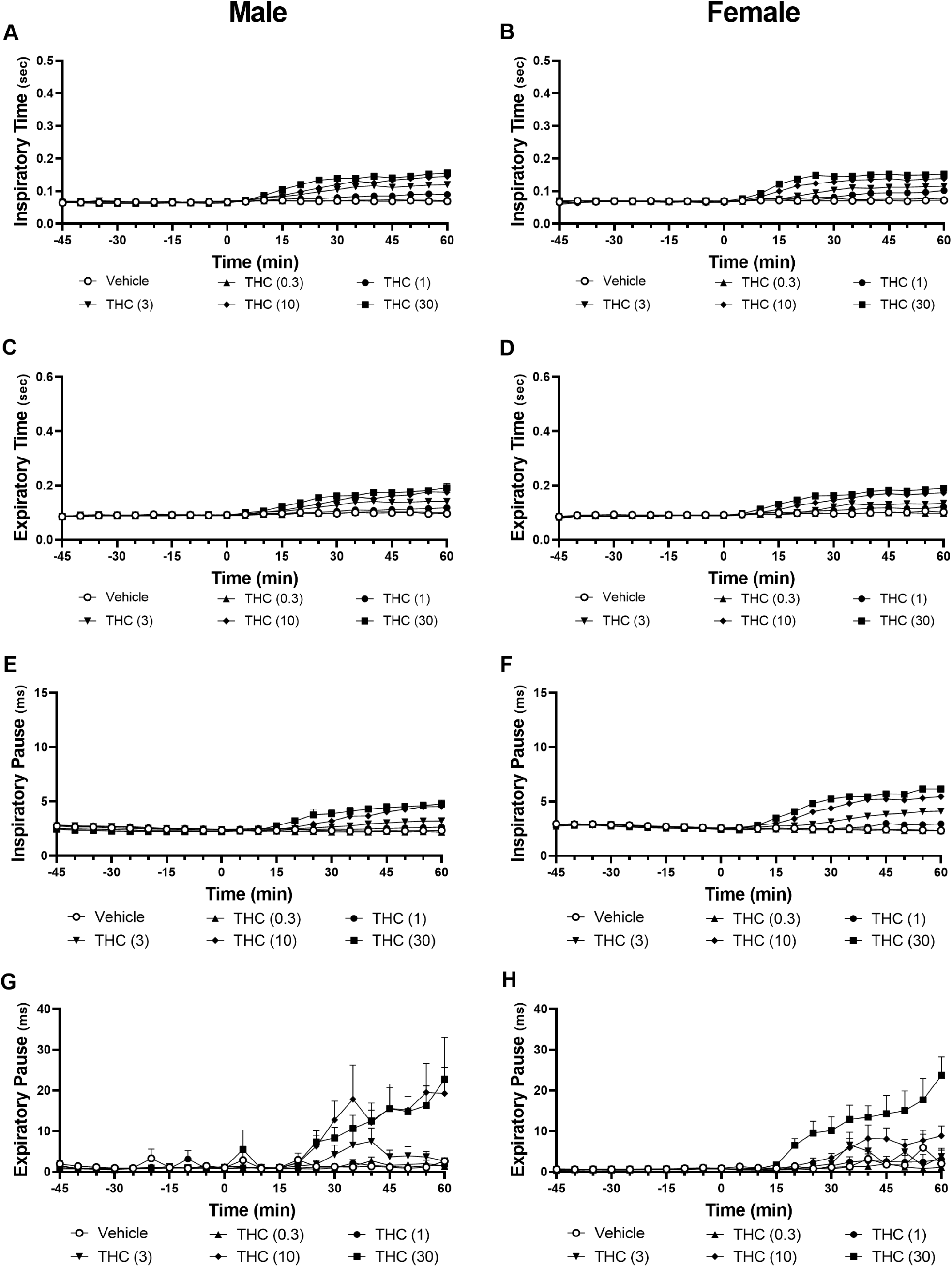
THC Inspiratory Time, Expiratory Time, End Inspiratory Pause, and End Expiratory Pause

### 3.3. Respiratory effects of THC and CP55,940 are reversed by AM251 and AM6545

Coadministration of AM251 (5 mg/kg) prevented the respiratory effects of CP55,940 (0.3 mg/kg) [Figure 11]. Animals that received both CP55,940 and AM251 exhibited respiratory parameters indistinguishable from vehicle controls [Table 5]. AM251 had similar effects on animals administered THC (10 mg/kg), attenuating all respiratory effects observed in animals administered THC alone [Figure 12; Table 6]. The respiratory parameters of animals that received both AM251 and THC demonstrated respiratory parameters did not differ from vehicle controls across all metrics.

**Figure 11:**
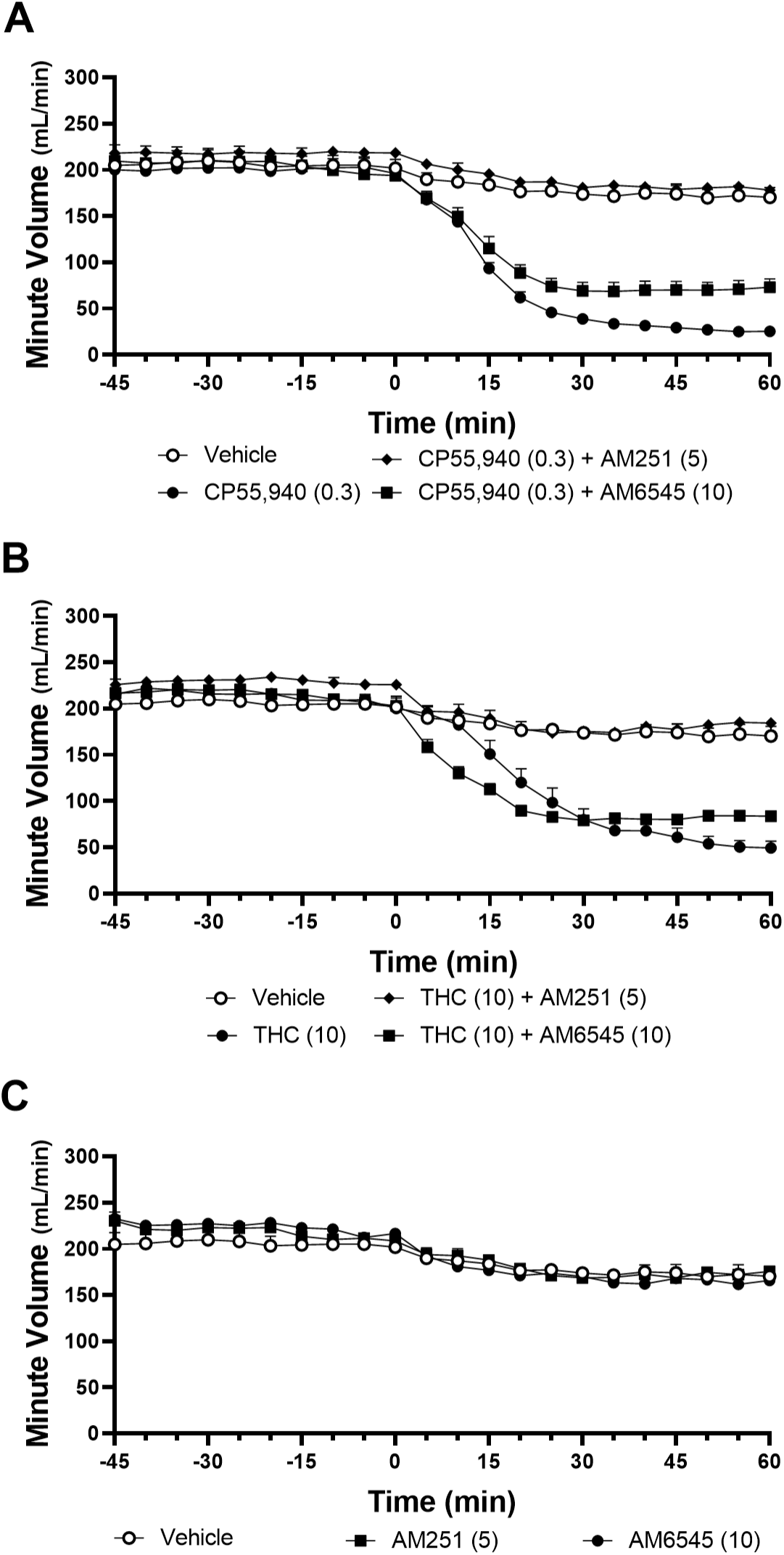
Antagonists Minute Ventilation and Antagonists Alone

**Figure 12:**
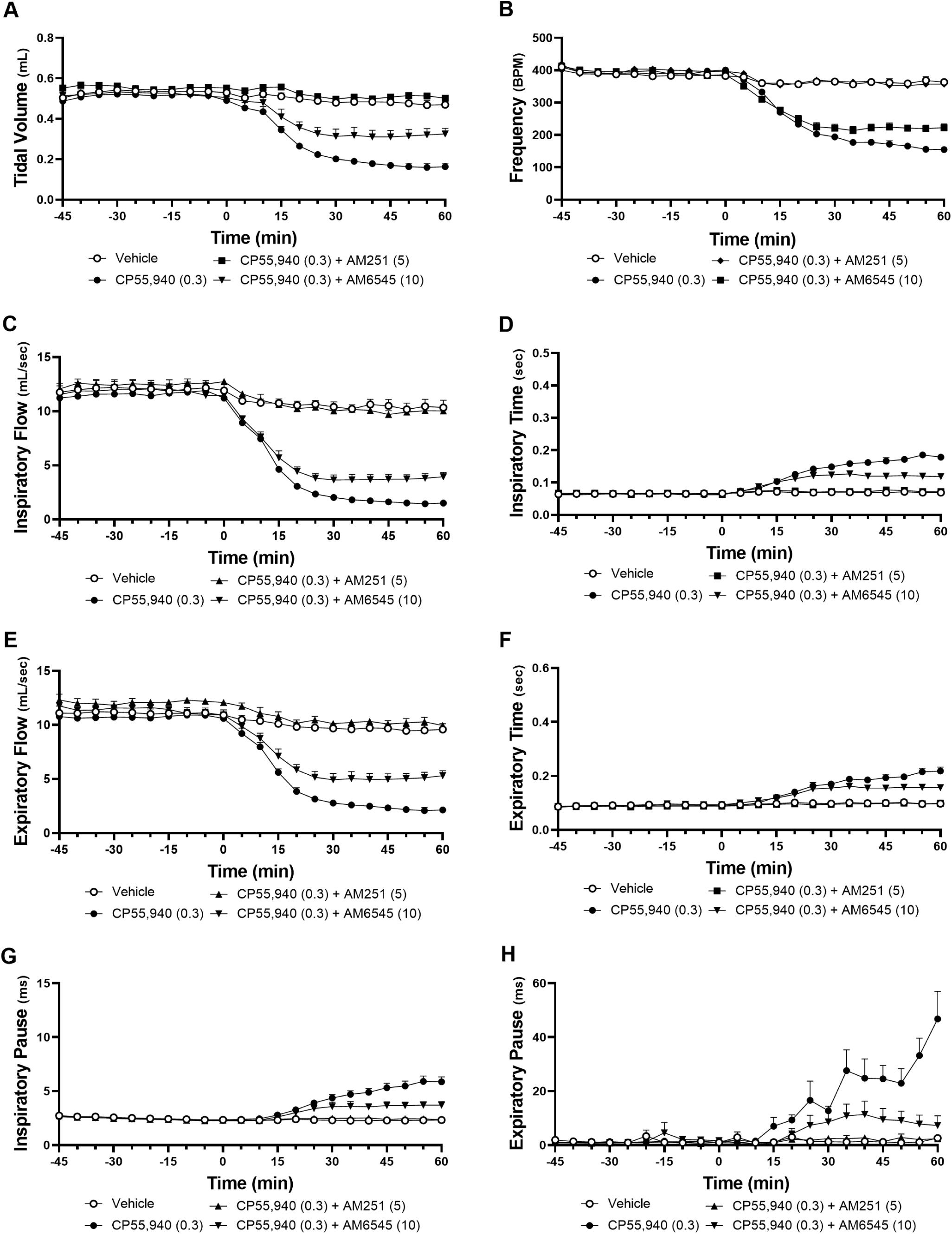
CP55,940 Antagonists non-minute Ventilation

**Table 5:** CP55,940 + AM251.

| Metric | Time |  | Drug |  | Interaction |  | vs.<br>CP55,940 | vs.<br>Vehicle |
| --- | --- | --- | --- | --- | --- | --- | --- | --- |
|  |  | p |  | p |  | p |  |  |
| Minute Ventilation | $F_{11,187} = 91.99$ | 0.0001 | $F_{2,17} = 161.7$ | 0.0001 | $F_{22,187} = 37.03$ | 0.0001 | 0.0001 | - |
| Tidal Volume | $F_{11,187} = 78.45$ | 0.0001 | $F_{2,17} = 99.65$ | 0.0001 | $F_{22,187} = 31.84$ | 0.0001 | 0.0001 | - |
| Inspiratory Flow | $F_{11,187} = 48.45$ | 0.0001 | $F_{2,17} = 153.6$ | 0.0001 | $F_{22,187} = 22.07$ | 0.0001 | 0.0001 | - |
| Expiratory Flow | $F_{11,187} = 73.08$ | 0.0001 | $F_{2,17} = 147.0$ | 0.0001 | $F_{22,187} = 28.17$ | 0.0001 | 0.0001 | - |
| Frequency | $F_{11,187} = 75.52$ | 0.0001 | $F_{2,17} = 127.2$ | 0.0001 | $F_{22,187} = 51.41$ | 0.0001 | 0.0001 | - |
| Inspiratory Time | $F_{11,187} = 52.91$ | 0.0001 | $F_{2,17} = 154.5$ | 0.0001 | $F_{22,187} = 50.51$ | 0.0001 | 0.0001 | - |
| Expiratory Time | $F_{11,187} = 43.52$ | 0.0001 | $F_{2,17} = 72.55$ | 0.0001 | $F_{22,187} = 33.44$ | 0.0001 | 0.0001 | - |
| Inspiratory Pause | $F_{11,187} = 37.18$ | 0.0001 | $F_{2,17} = 68.03$ | 0.0001 | $F_{22,187} = 34.43$ | 0.0001 | 0.0001 | - |
| Expiratory Pause | $F_{11,187} = 9.497$ | 0.0001 | $F_{2,17} = 35.45$ | 0.0001 | $F_{22,187} = 8.598$ | 0.0001 | 0.0001 | - |

**Table 6:** THC + AM251.

| Metric | Time |  | Drug |  | Interaction |  | vs.<br>THC | vs.<br>Vehicle |
| --- | --- | --- | --- | --- | --- | --- | --- | --- |
|  |  | p |  | p |  | p |  |  |
| Minute Ventilation | $F_{11,187} = 91.99$ | 0.0001 | $F_{2,17} = 161.7$ | 0.0001 | $F_{22,187} = 37.03$ | 0.0001 | 0.0001 | - |
| Tidal Volume | $F_{11,187} = 78.45$ | 0.0001 | $F_{2,17} = 99.65$ | 0.0015 | $F_{22,187} = 31.84$ | 0.0001 | 0.0001 | - |
| Inspiratory Flow | $F_{11,187} = 48.45$ | 0.0001 | $F_{2,17} = 153.6$ | 0.0001 | $F_{22,187} = 22.07$ | 0.0001 | 0.0001 | - |
| Expiratory Flow | $F_{11,187} = 73.08$ | 0.0001 | $F_{2,17} = 147.0$ | 0.0001 | $F_{22,187} = 28.17$ | 0.0001 | 0.0001 | - |
| Frequency | $F_{11,187} = 75.52$ | 0.0001 | $F_{2,17} = 127.2$ | 0.0001 | $F_{22,187} = 51.41$ | 0.0001 | 0.0001 | - |
| Inspiratory Time | $F_{11,187} = 54.56$ | 0.0001 | $F_{2,17} = 138.0$ | 0.0001 | $F_{22,187} = 51.85$ | 0.0001 | 0.0001 | - |
| Expiratory Time | $F_{11,187} = 43.52$ | 0.0001 | $F_{2,17} = 72.55$ | 0.0001 | $F_{22,187} = 33.44$ | 0.0001 | 0.0001 | - |
| Inspiratory Pause | $F_{11,187} = 37.18$ | 0.0001 | $F_{2,17} = 68.03$ | 0.0001 | $F_{22,187} = 34.43$ | 0.0001 | 0.0001 | - |
| Expiratory Pause | $F_{11,187} = 9.497$ | 0.0561 | $F_{2,17} = 35.45$ | 0.0001 | $F_{22,187} = 8.598$ | 0.0001 | 0.0001 | - |

AM6545 lessened the severity of the respiratory effects of CP55,940, though much more modestly than AM251 [Figure 11; Table 7]. Most metrics demonstrated lesser suppression than in animals receiving only CP55,940, apart from expiratory time, which did not differ from animals who only received CP55,940 [Figure 13]. However, these metrics failed to normalize to vehicle levels, apart from end expiratory pause, which did not differ from vehicle animals. Similarly, AM6545 broadly failed to antagonize the respiratory suppressive effects of THC, as overall minute ventilation, frequency, tidal volume, inspiratory flow, expiratory flow, inspiratory time, and expiratory time flow did not differ from animals administered only THC [Table 8]. Animals that received AM6545 did show improvements to inspiratory and expiratory pause in comparison to animals that received only THC, though inspiratory pause failed to normalize to vehicle levels. Neither AM251 nor AM6545 affected respiratory parameters when administered alone (Supplemental 1).

**Figure 13:**
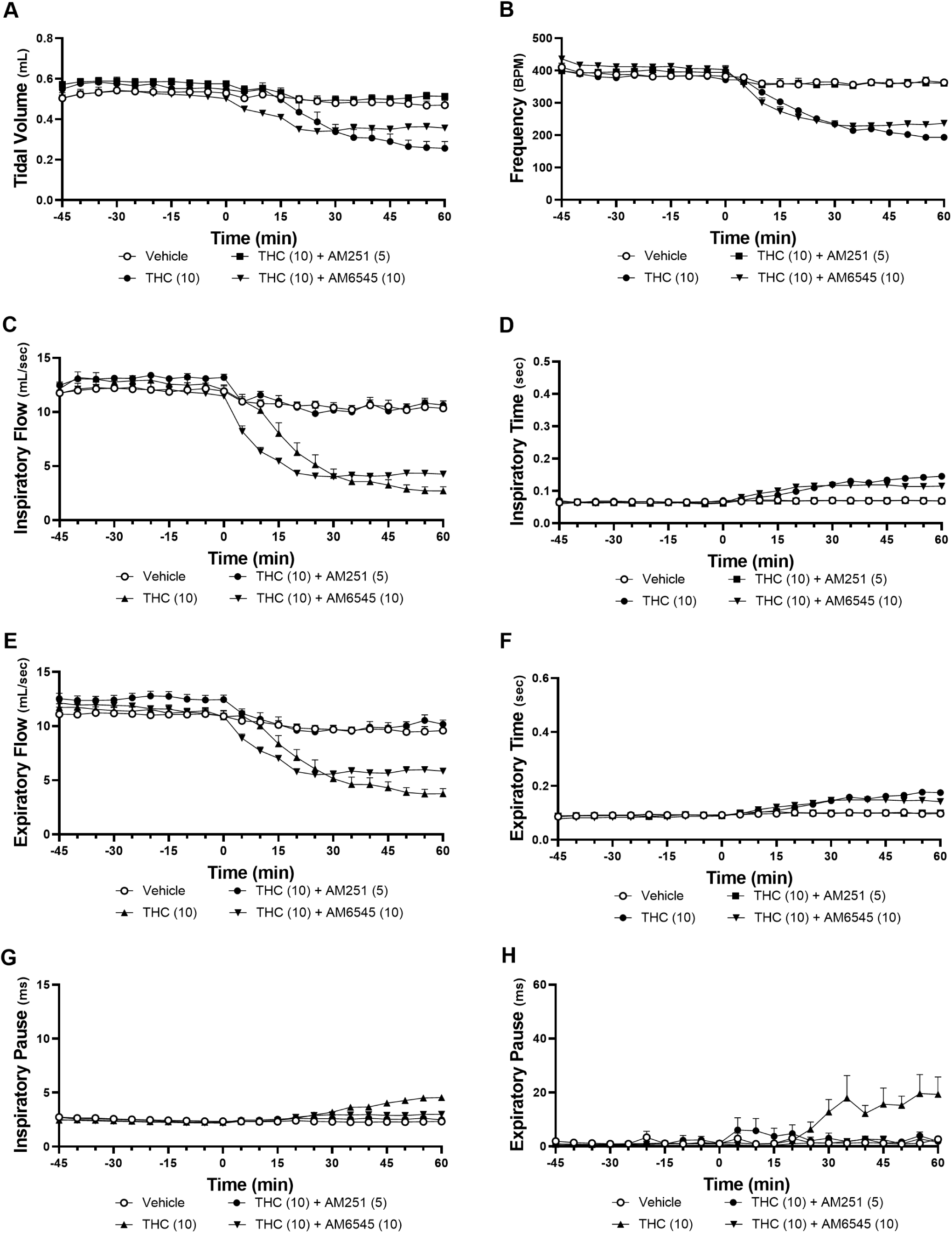
THC Antagonists non-Minute Ventilation

**Table 7:**
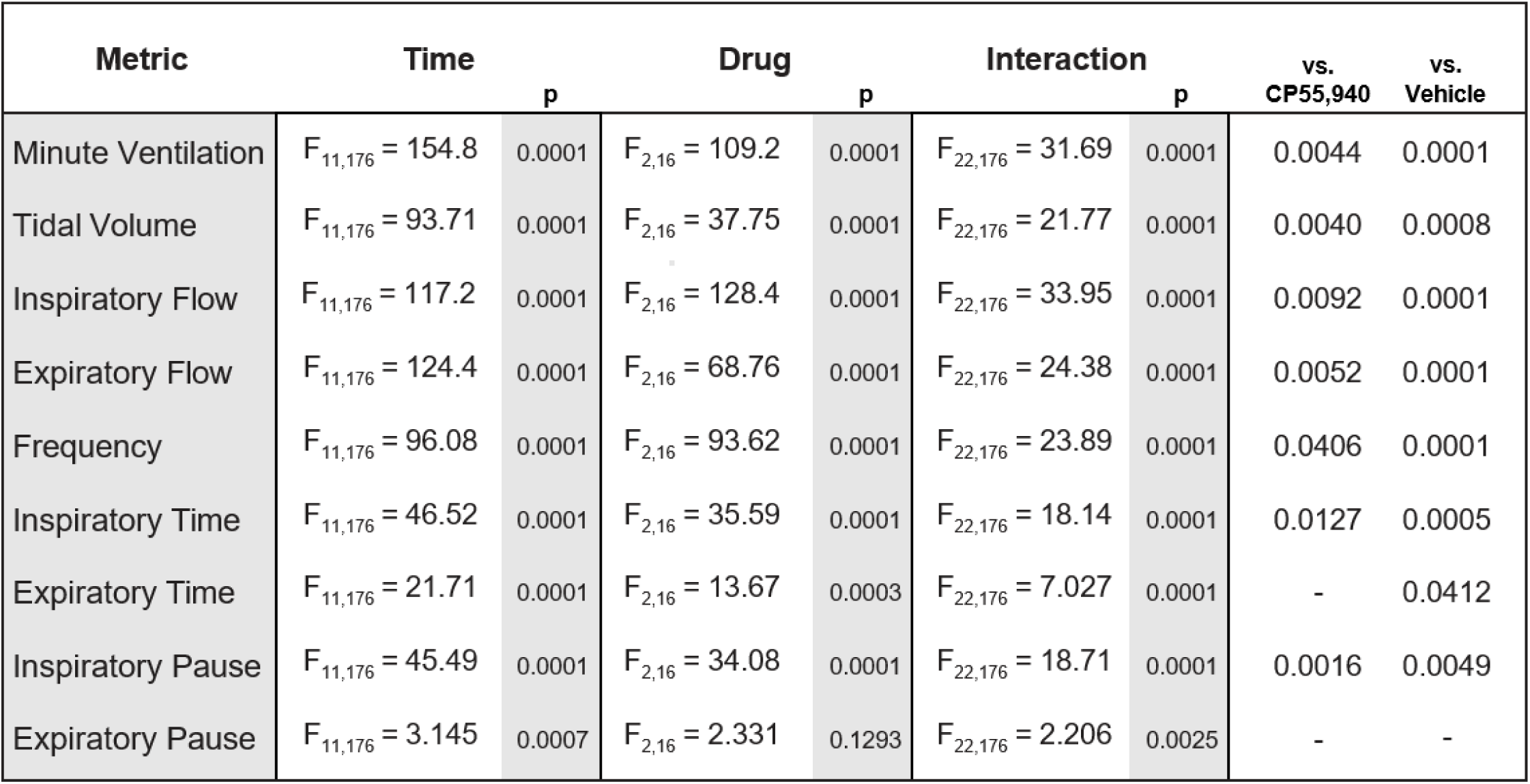
CP55,940 + AM6545.

| Metric | Time |  | Drug |  | Interaction |  | vs.<br>CP55,940 | vs.<br>Vehicle |
| --- | --- | --- | --- | --- | --- | --- | --- | --- |
|  |  | p |  | p |  | p |  |  |
| Minute Ventilation | $F_{11,176} = 154.8$ | 0.0001 | $F_{2,16} = 109.2$ | 0.0001 | $F_{22,176} = 31.69$ | 0.0001 | 0.0044 | 0.0001 |
| Tidal Volume | $F_{11,176} = 93.71$ | 0.0001 | $F_{2,16} = 37.75$ | 0.0001 | $F_{22,176} = 21.77$ | 0.0001 | 0.0040 | 0.0008 |
| Inspiratory Flow | $F_{11,176} = 117.2$ | 0.0001 | $F_{2,16} = 128.4$ | 0.0001 | $F_{22,176} = 33.95$ | 0.0001 | 0.0092 | 0.0001 |
| Expiratory Flow | $F_{11,176} = 124.4$ | 0.0001 | $F_{2,16} = 68.76$ | 0.0001 | $F_{22,176} = 24.38$ | 0.0001 | 0.0052 | 0.0001 |
| Frequency | $F_{11,176} = 96.08$ | 0.0001 | $F_{2,16} = 93.62$ | 0.0001 | $F_{22,176} = 23.89$ | 0.0001 | 0.0406 | 0.0001 |
| Inspiratory Time | $F_{11,176} = 46.52$ | 0.0001 | $F_{2,16} = 35.59$ | 0.0001 | $F_{22,176} = 18.14$ | 0.0001 | 0.0127 | 0.0005 |
| Expiratory Time | $F_{11,176} = 21.71$ | 0.0001 | $F_{2,16} = 13.67$ | 0.0003 | $F_{22,176} = 7.027$ | 0.0001 | - | 0.0412 |
| Inspiratory Pause | $F_{11,176} = 45.49$ | 0.0001 | $F_{2,16} = 34.08$ | 0.0001 | $F_{22,176} = 18.71$ | 0.0001 | 0.0016 | 0.0049 |
| Expiratory Pause | $F_{11,176} = 3.145$ | 0.0007 | $F_{2,16} = 2.331$ | 0.1293 | $F_{22,176} = 2.206$ | 0.0025 | - | - |

**Table 8:** THC + AM6545.

| Metric | Time |  | Drug |  | Interaction |  | vs.<br>THC | vs.<br>Vehicle |
| --- | --- | --- | --- | --- | --- | --- | --- | --- |
|  |  | p |  | p |  | p |  |  |
| Minute Ventilation | $F_{11,176} = 167.6$ | 0.0001 | $F_{2,16} = 42.19$ | 0.0001 | $F_{22,176} = 48.68$ | 0.0001 | - | 0.0001 |
| Tidal Volume | $F_{11,176} = 69.74$ | 0.0001 | $F_{2,16} = 8.666$ | 0.0028 | $F_{22,176} = 29.06$ | 0.0001 | - | 0.0088 |
| Inspiratory Flow | $F_{11,176} = 109.1$ | 0.0001 | $F_{2,16} = 46.69$ | 0.0001 | $F_{22,176} = 47.88$ | 0.0001 | - | 0.0001 |
| Expiratory Flow | $F_{11,176} = 104.6$ | 0.0001 | $F_{2,16} = 29.40$ | 0.0001 | $F_{22,176} = 31.69$ | 0.0001 | - | 0.0001 |
| Frequency | $F_{11,176} = 115.7$ | 0.0001 | $F_{2,16} = 94.31$ | 0.0001 | $F_{22,176} = 28.16$ | 0.0001 | - | 0.0001 |
| Inspiratory Time | $F_{11,176} = 95.01$ | 0.0001 | $F_{2,16} = 62.91$ | 0.0001 | $F_{22,176} = 42.39$ | 0.0001 | - | 0.0001 |
| Expiratory Time | $F_{11,176} = 59.91$ | 0.0001 | $F_{2,16} = 27.54$ | 0.0001 | $F_{22,176} = 15.58$ | 0.0001 | - | 0.0001 |
| Inspiratory Pause | $F_{11,176} = 34.83$ | 0.0001 | $F_{2,16} = 37.22$ | 0.0001 | $F_{22,176} = 17.87$ | 0.0001 | 0.0026 | 0.0014 |
| Expiratory Pause | $F_{11,176} = 6.260$ | 0.0001 | $F_{2,16} = 8.835$ | 0.0026 | $F_{22,176} = 3.681$ | 0.0001 | 0.0057 | - |

### 3.4. CB_1_ mRNA is expressed in the pre-Bötzinger Complex

RNAScope *in situ* hybridization confirmed the presence of CB_1_ mRNA in the preBötC [Figure 14]. Labeling was sparse but consistent in wild type animals. CB_1_ mRNA was detected in 27.6% of preBötC neurons. 25.2% of CB_1_-expressing cells also expressed VGLUT2 mRNA, making up 28.2% of the total population of observed VGLUT2-expressing cells. In knockout animals, minimal CB_1_ staining was observed; only 1.7% of preBötC neurons expressed CB_1_ mRNA [Figure 14a].

**Figure 14:**
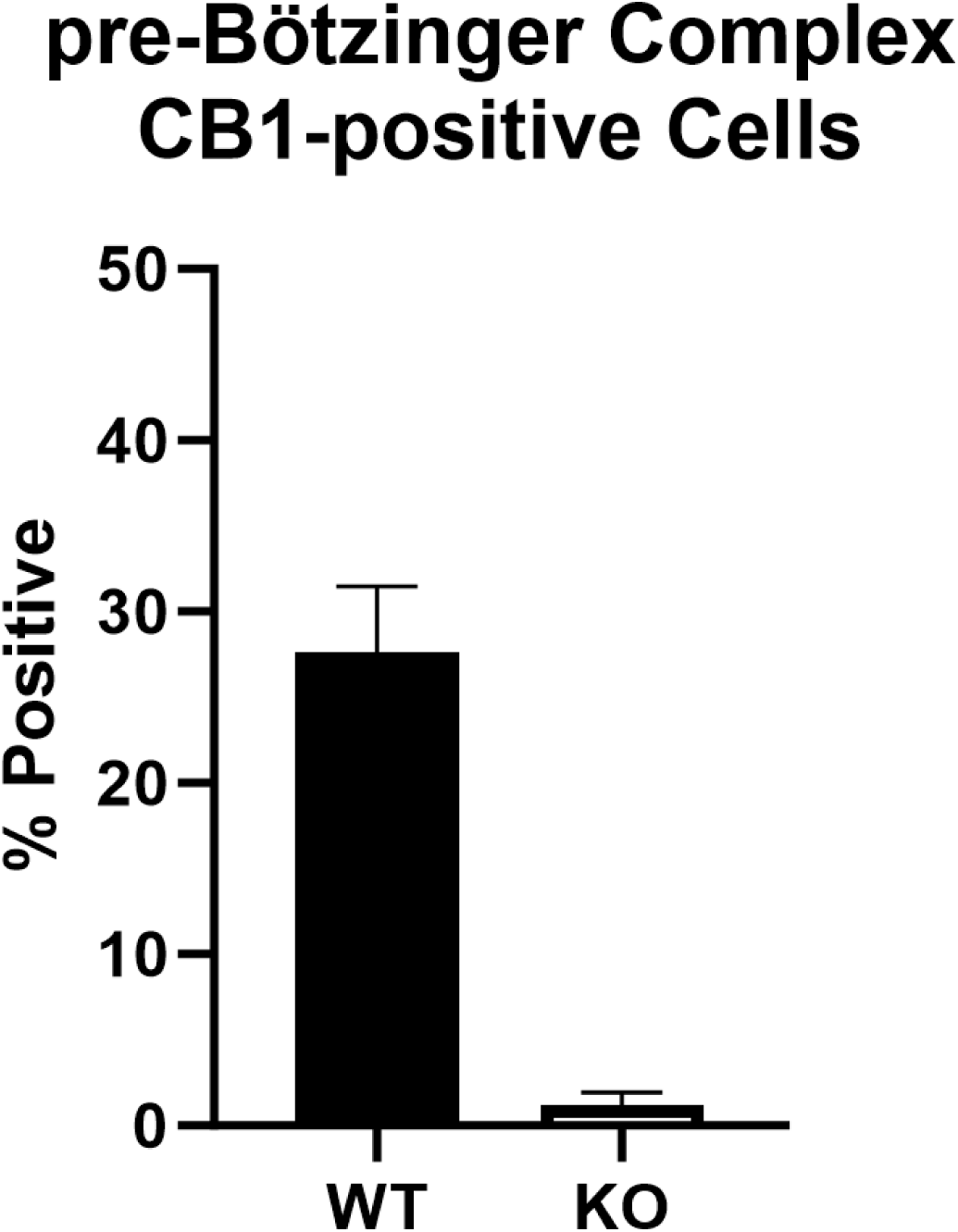
RNAScope *in situ* hybridization 14a)

**Figure 14b).**
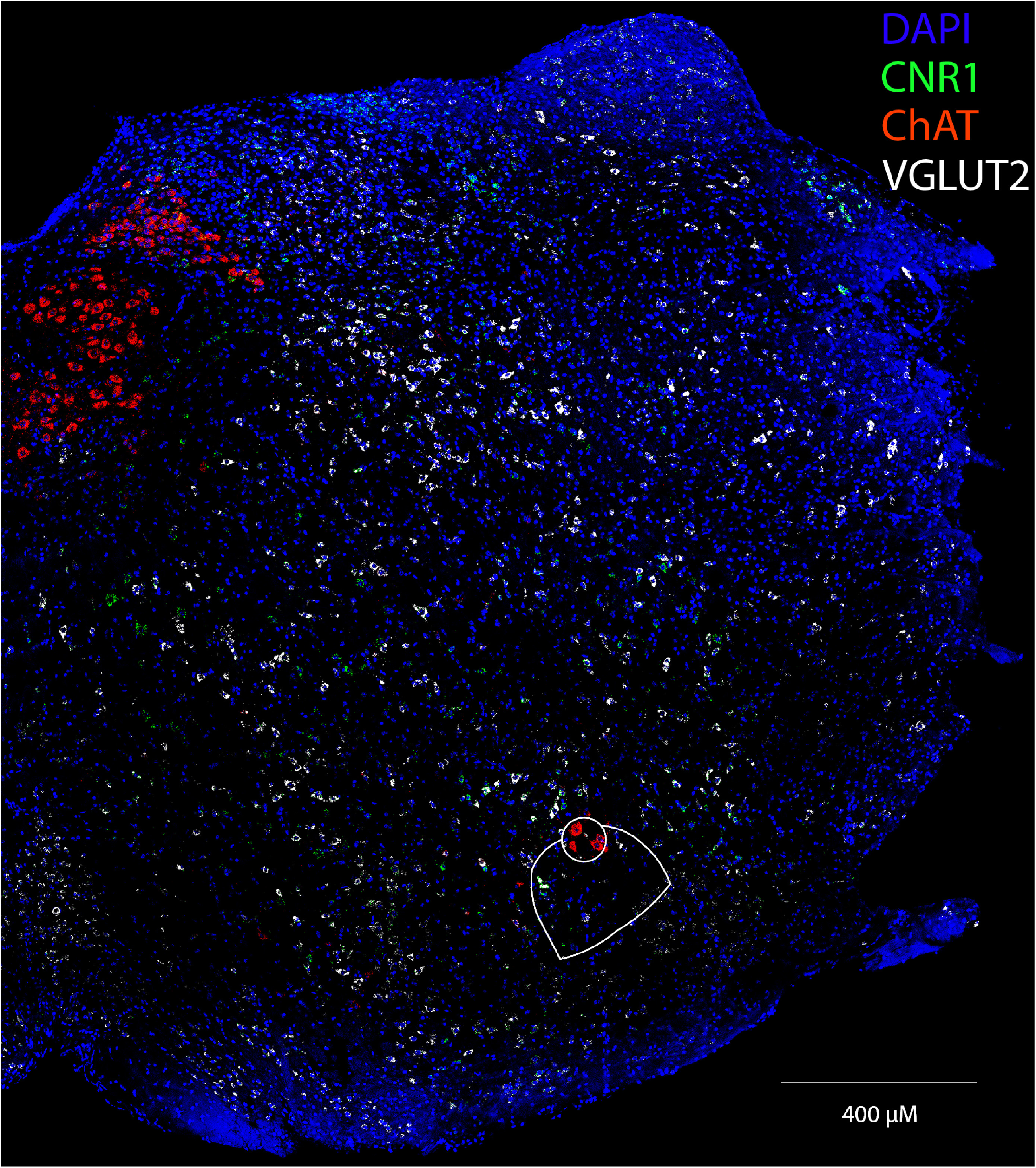
Confocal image (20x) of a wild type mouse coronal slice at the level of the pre-Bötzinger Complex. CB1 mRNA is labeled in green, VGLUT2 mRNA is labeled in white, and ChAT mRNA (used as a marker for the nucleus ambiguous) is labeled in red. Nucleus ambiguous (top) and pre-Botzinger Complex (bottom) outlined.

**Figure 14c).**
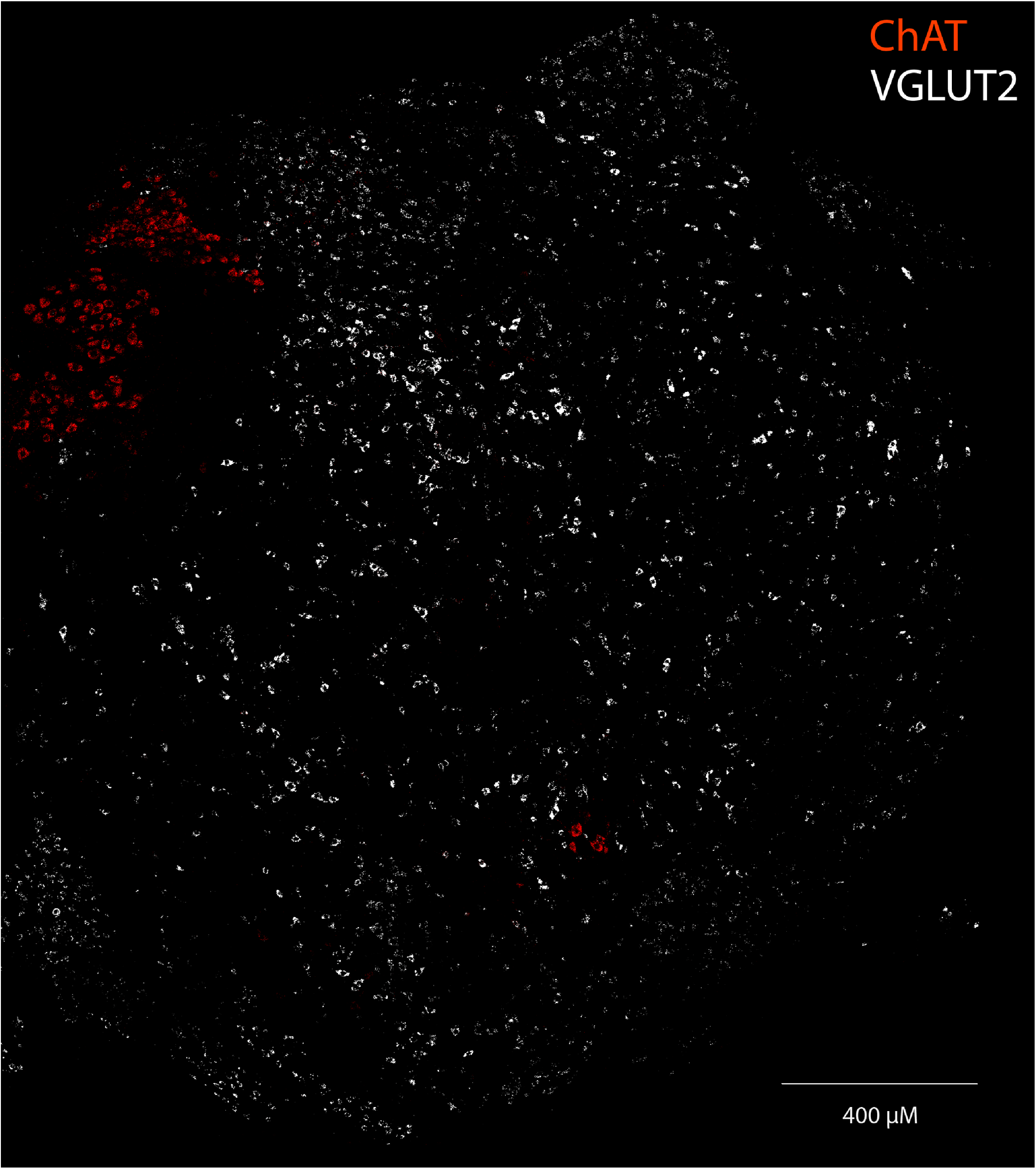
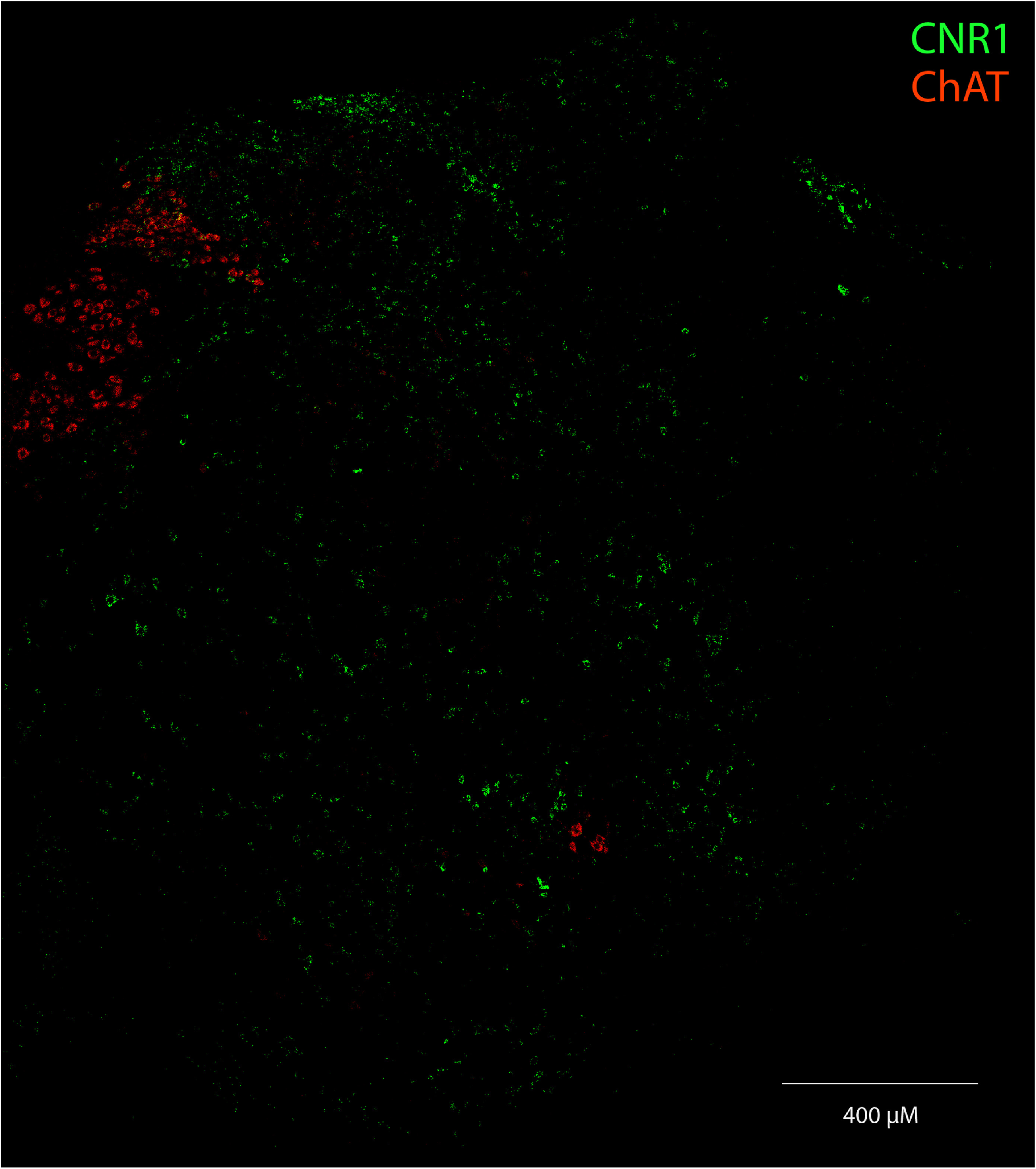
The same medullary slice displaying only the ChAT and VGLUT2 channels (top) and ChAT and CB1 channels (bottom)

**Figure 14d).**
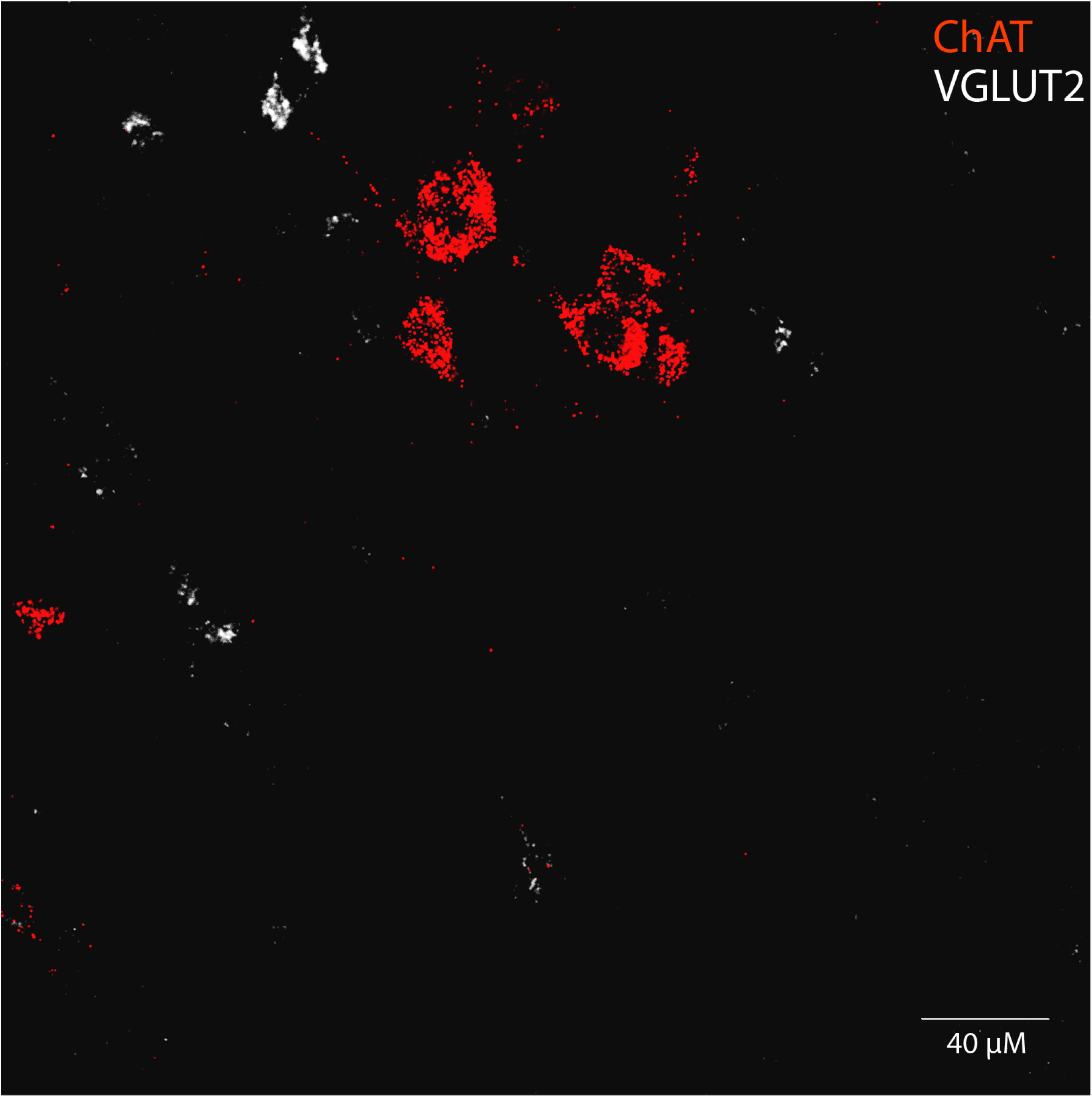
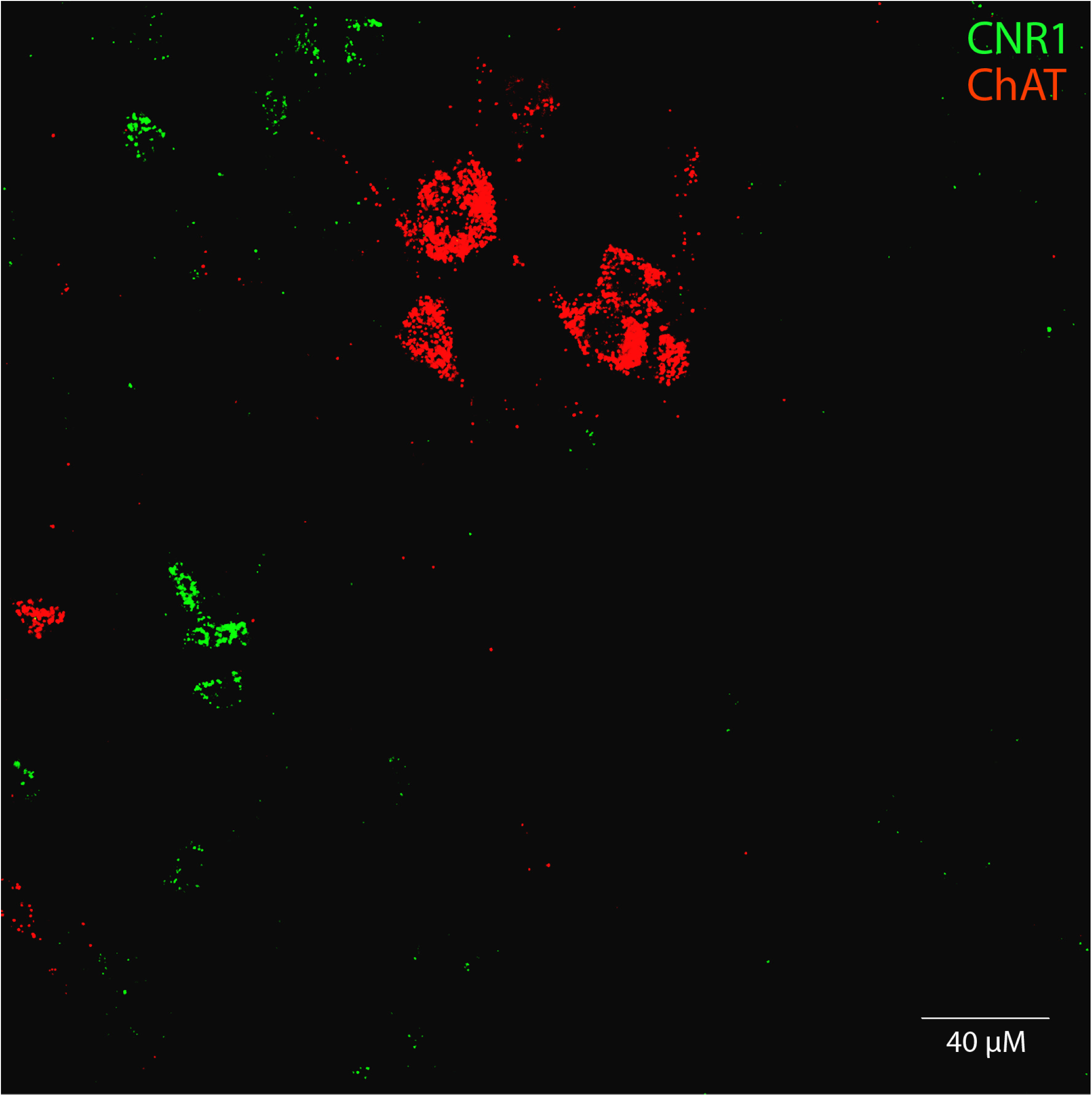
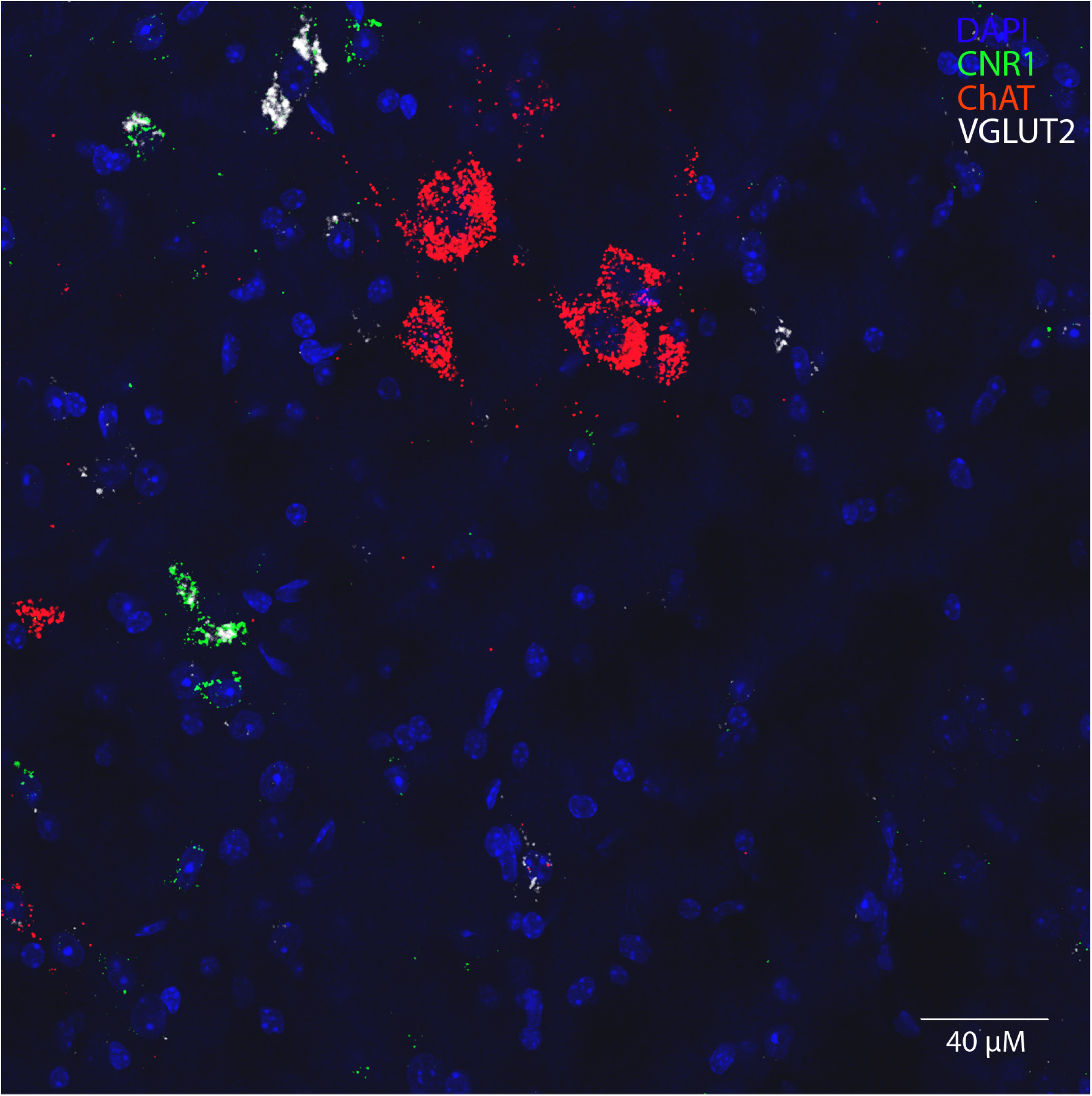
The same slice at higher resolution. ChAT and VGLUT2 signals (top), ChAT and CB1 signals (middle), and all channels merged (bottom)

**Figure 14e).**
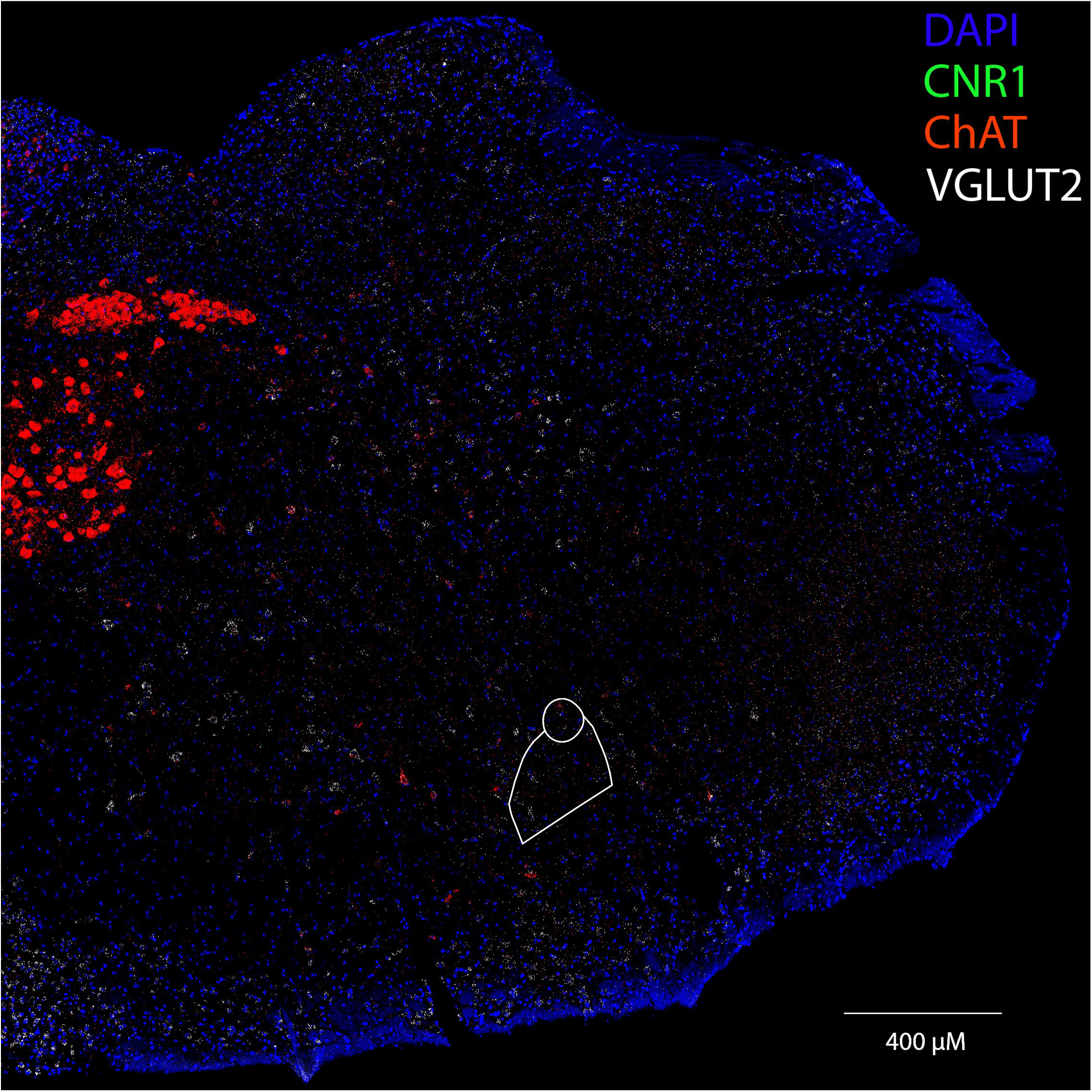
Confocal image (20x) of a CB_1_^−/−^ mouse coronal slice at the level of the pre-Bötzinger Complex. CB1 mRNA is labeled in green, VGLUT2 mRNA is labeled in white, and ChAT mRNA (used as a marker for the nucleus ambiguous) is labeled in red. Nucleus ambiguous (top) and pre-Botzinger Complex (bottom) outlined.

**Figure 14f).**
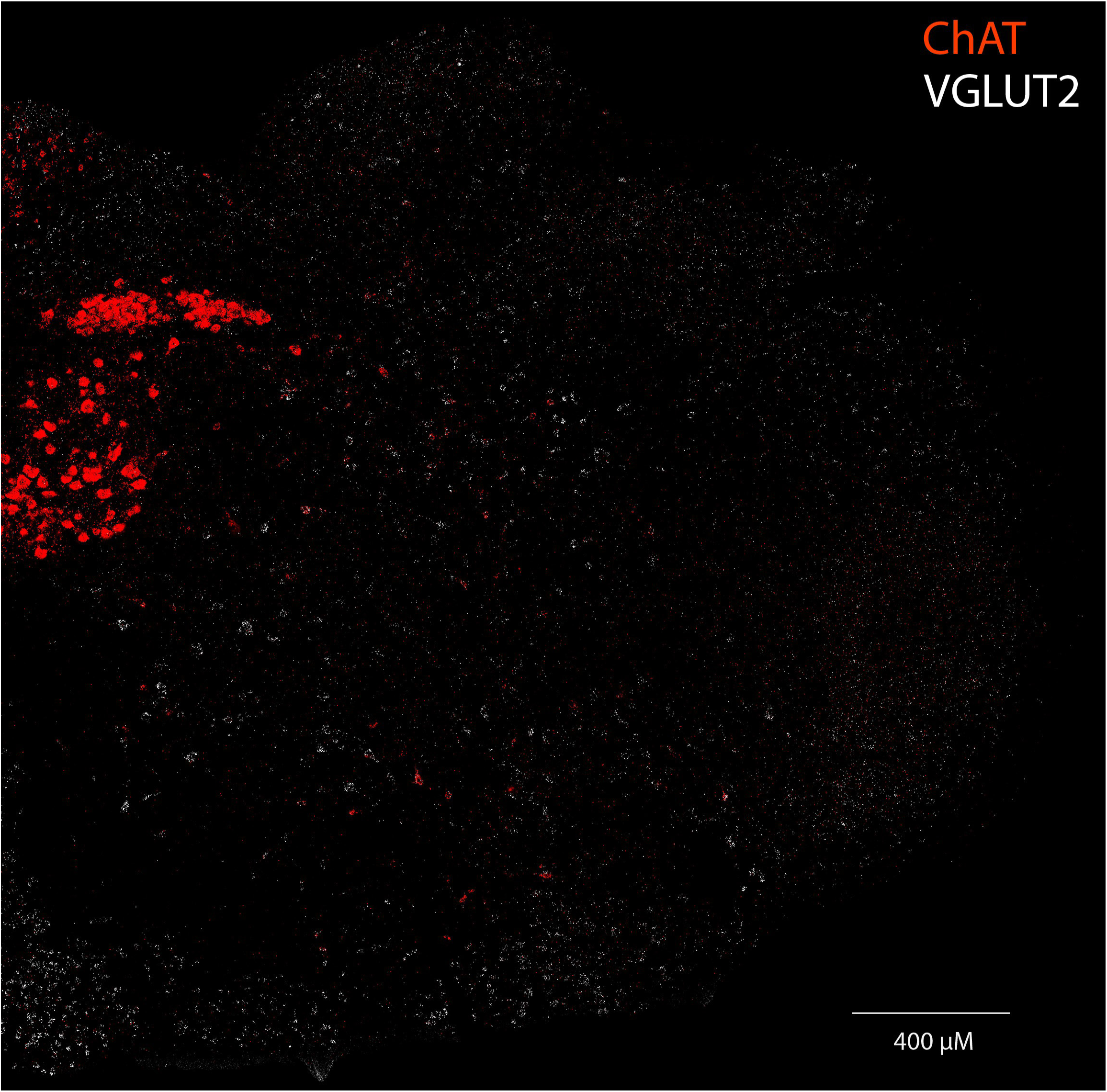
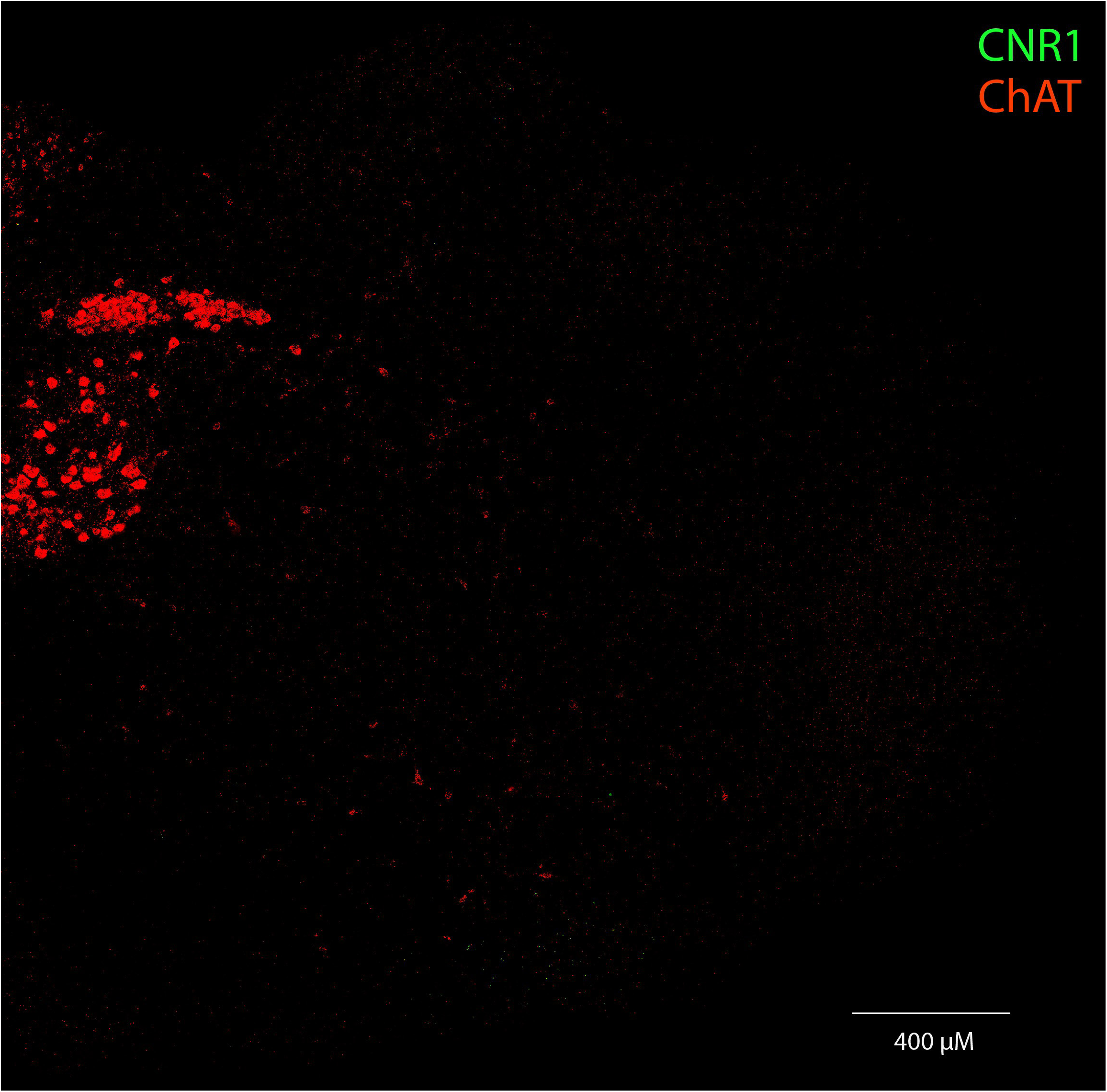
The same slice displaying only the ChAT and VGLUT2 channels (top) and ChAT and CB1 channels (bottom).

**Figure 14g).**
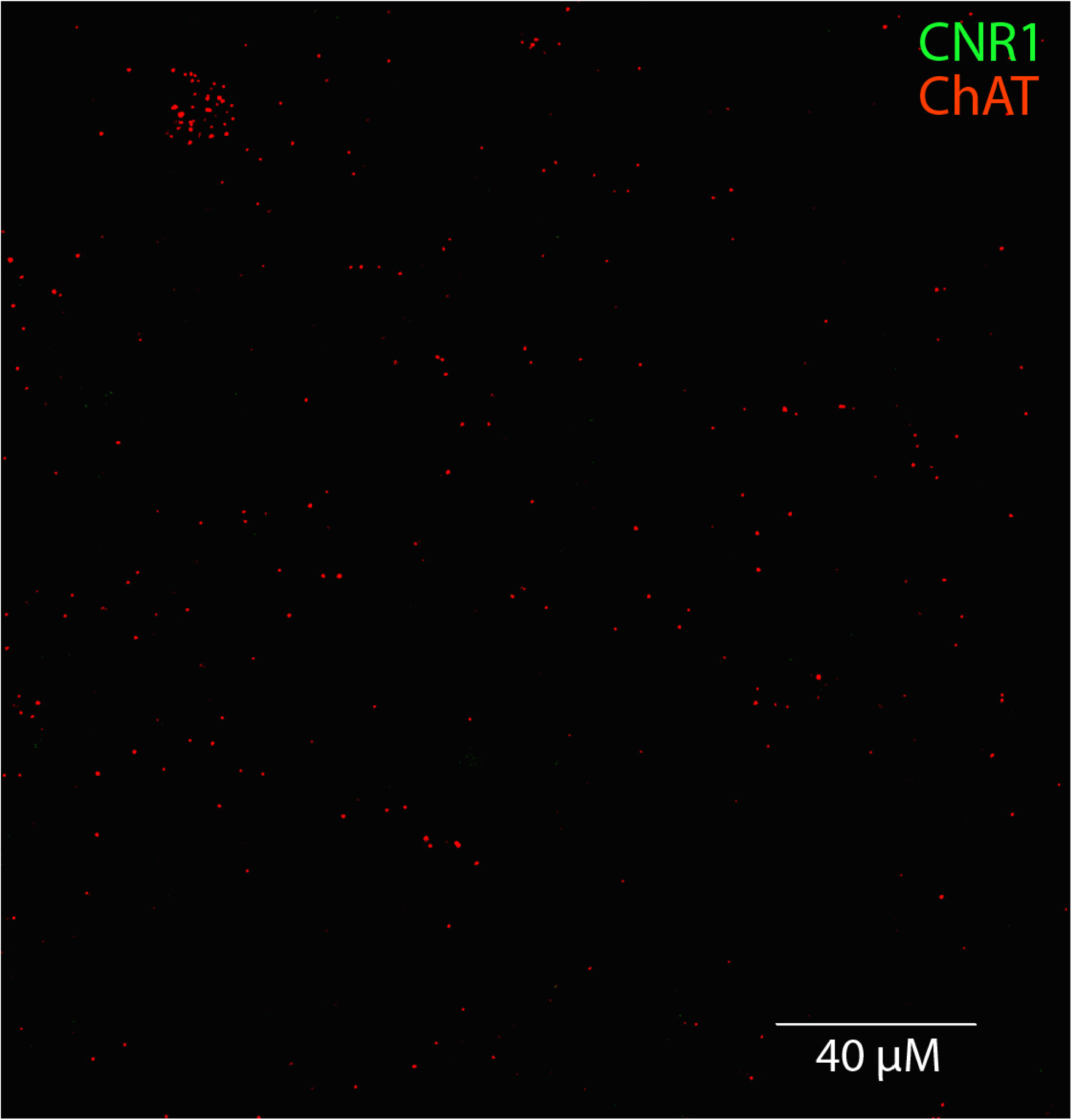
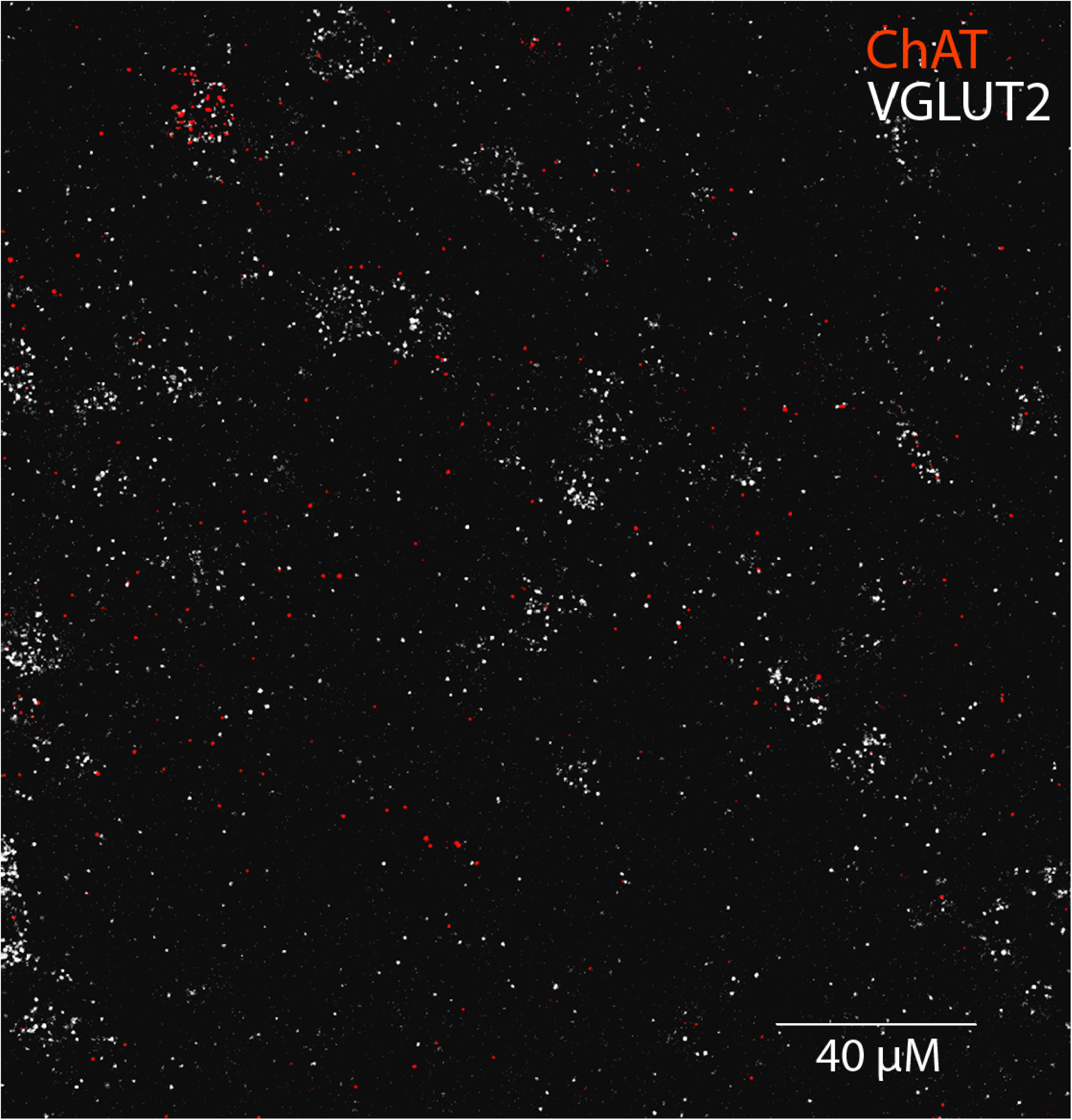
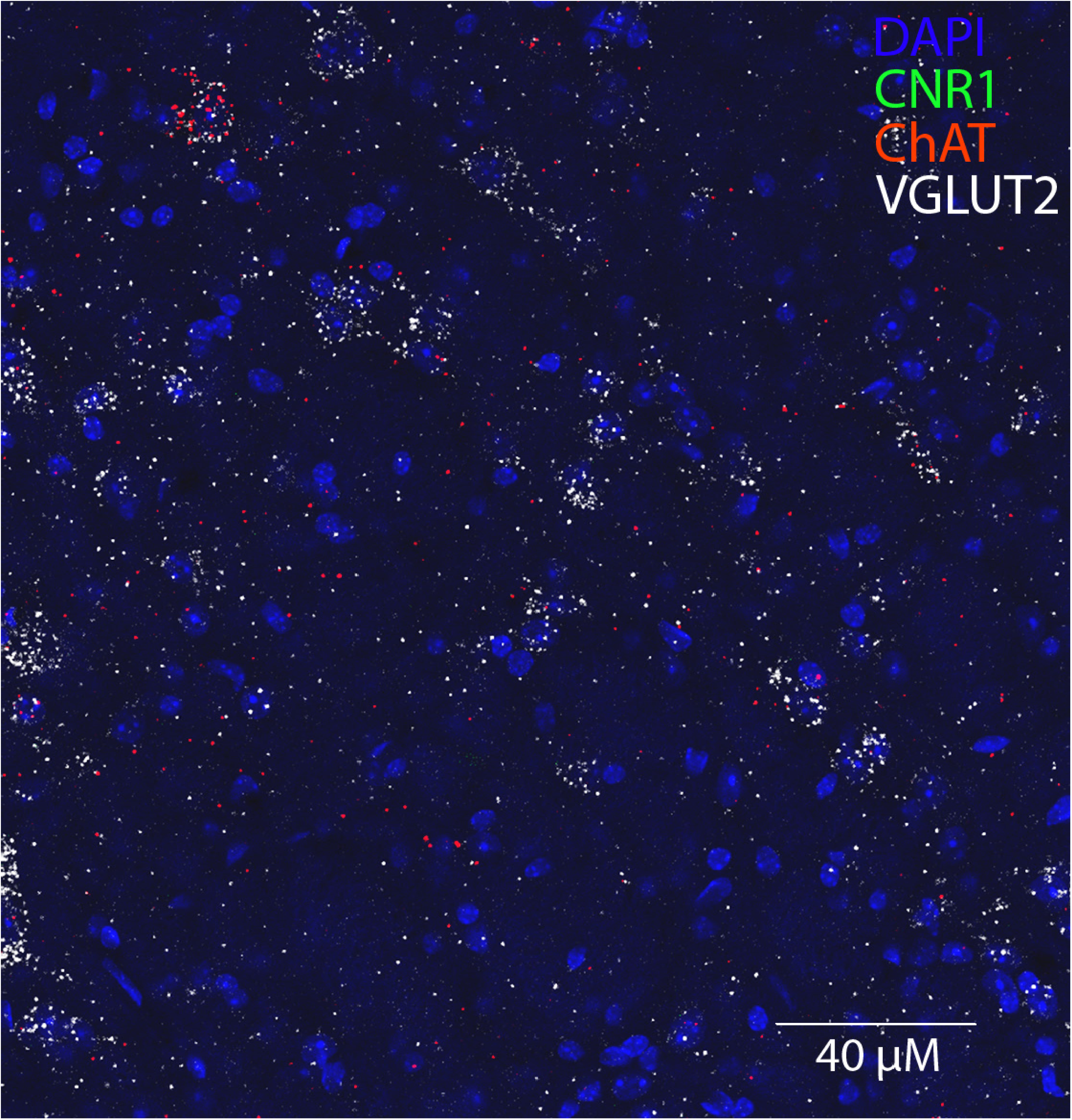
The same slice at higher resolution. ChAT and VGLUT2 signals (top), ChAT and CB1 signals (middle), and all channels merged (bottom)

**Figure 14l).**
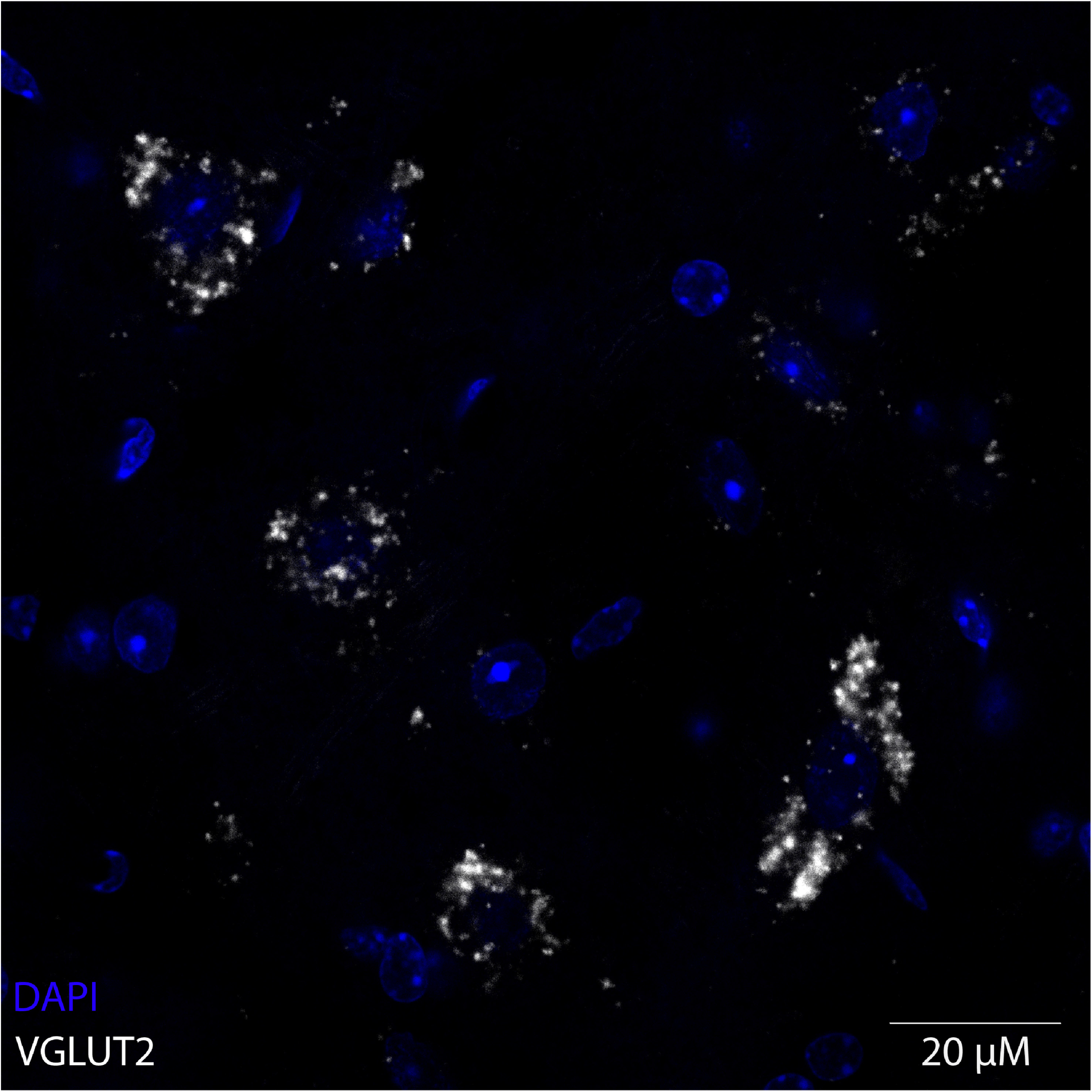
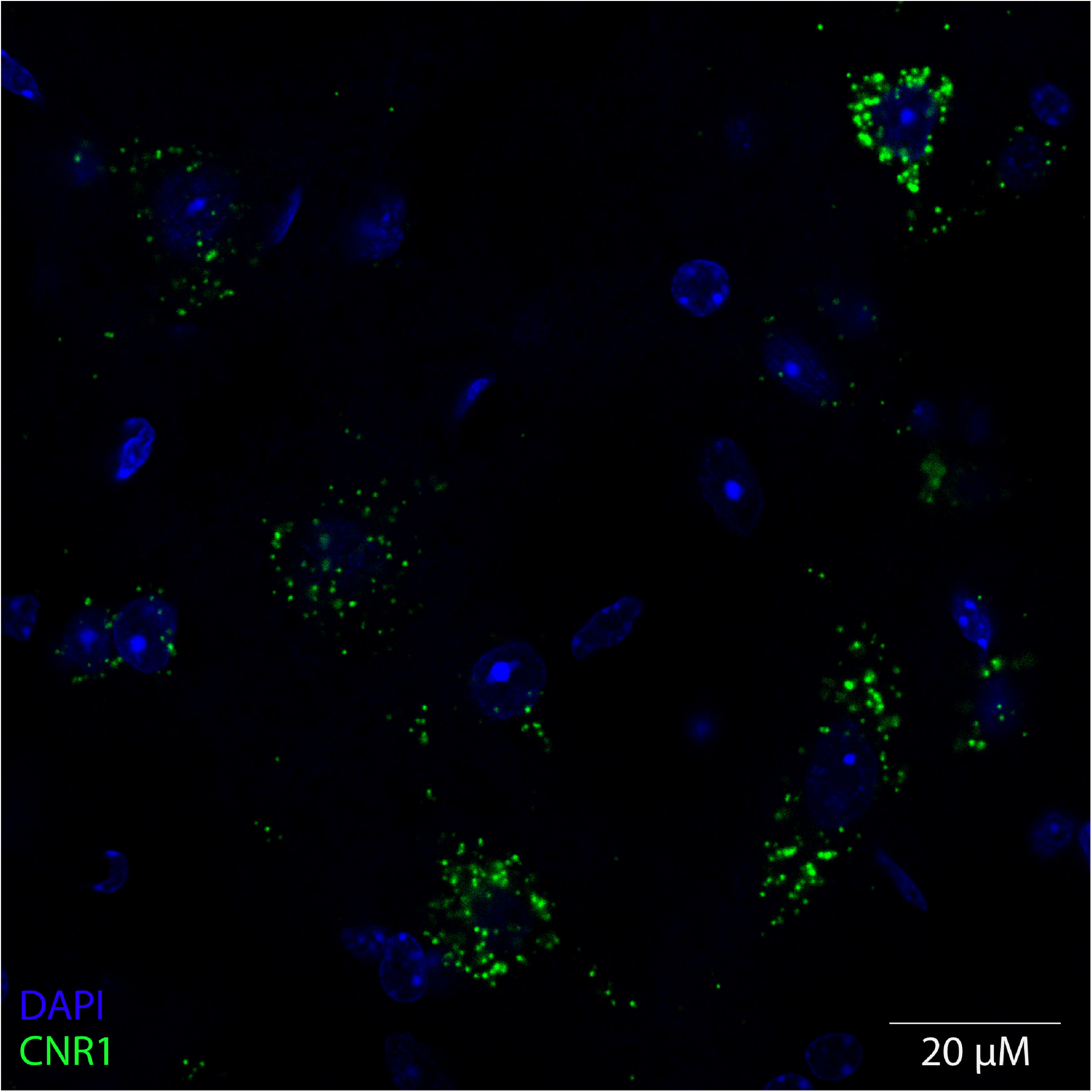
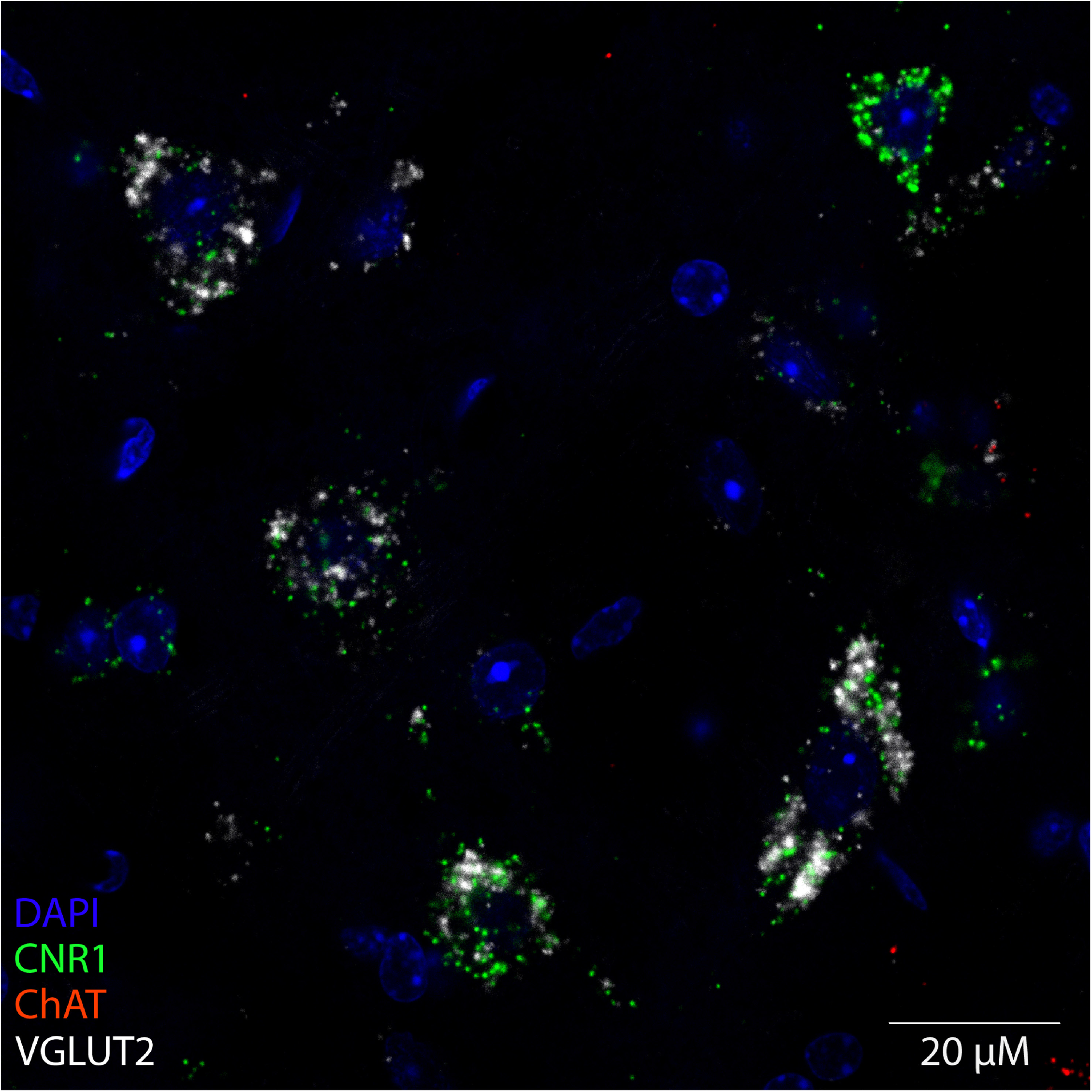
Confocal 60x image of medullary neurons coexpressing VGLUT2 (top) and CB_1_ (middle) mRNA. Merged image (bottom).

## 4. Discussion

Users of illicit synthetic cannabinoids suffer from a variety of life-threatening side effects, including cardiovascular complications and acute respiratory depression [15–21]. Virtually nothing is known about how CB_1_ activation alters respiratory function, and existing studies examining cannabinoid interactions with the respiratory system primarily focus on targeting other members of the endocannabinoid system (e.g., the Cannabinoid Type 2 receptor) to ameliorate the effects of opioid drugs [28, 29]. Early neuroanatomical studies supported the idea that cannabinoid agonists would have minimal effect on the respiratory system; binding and autoradiography studies revealed that expression of cannabinoid receptors in the medulla, where breathing is primarily controlled, were very low in comparison to regions where cannabinoid receptors are densely expressed (e.g., basal ganglia, cerebellum, hippocampus, cortex), explaining why even high doses of cannabis are rarely lethal [10]. However, clinical reports of synthetic cannabinoid overdose involving respiratory depression suggest that the respiratory effects of synthetic cannabinoids targeting CB_1_ must be considered.

In recognition of the danger posed by these compounds, various attempts to regulate synthetic cannabinoids have been implemented; in the United States several synthetic cannabinoids were federally scheduled in 2011, and more have been periodically added as they have been identified in human use [30]. Most recently to the time of publication, six synthetic cannabinoids were provisionally classified as Schedule 1 in December of 2023 [31]. However, use has persisted despite the federal ban; in 2022, 3.2% of US high school seniors reported synthetic cannabinoid use [32].

Synthetic cannabinoid use and overdose rates are poorly tracked. A variety of factors make accurate prevalence difficult to ascertain, including unstable pharmacology, a lack of consistent testing, and the absence of standardized, commercially available tests. Despite these factors, synthetic cannabinoids remain the largest and most widely used class of new psychoactive compounds in the United States, and coadministration of synthetic cannabinoids and opioids has recently been documented in lethal overdose [22, 23, 33]. No antidote is available for synthetic cannabinoid overdose, and treatment is largely limited to supportive care [17]. Attempts to treat cannabinoid-induced respiratory depression with existing reversal agents (e.g., Naloxone) have met with generally negative results [34–36].

Here we used whole body plethysmography to definitively demonstrate that both the synthetic bicyclic cannabinoid CP55,940 and the phytocannabinoid THC can induce respiratory depression. CP55,940 and THC dose-dependently attenuated respiratory function to a striking degree. Observed changes were rapid, persistent, and observed in all tested metrics, impacting each phase of the respiratory cycle by reducing inspiratory and expiratory flow, as well as extending inspiratory and expiratory time. These effects translated into severe depression of respiratory frequency, tidal volume, and minute ventilation that persisted for over one hour.

While all parameters were impacted by at least one tested dose of both CP55,940 and THC, the doses at which these effects were observed were not uniform. Reductions to parameters that measure the characteristics of individual breaths (i.e., flow rate and phase time) consistently developed before effects on parameters that measure the time between respiratory phases (i.e., inspiratory and expiratory pause), suggesting that changes to respiratory frequency and amplitude were primarily mediated by changes to the characteristics of individual breaths rather than by changes to the interval between them. Sustained cessations of breathing (apnea), as suggested by significant elongation of the post-expiratory pause, were observed with high doses of both drugs, indicating that cannabinoid agonists can induce the periods of sustained apnea that impart lethality to drugs that target the mu opioid receptor. This supports the hypothesis that cannabinoid agonists are acting on the respiratory system at sites of action that overlap with those affected by opioids, though further work is required to determine whether CB_1_ receptors are coexpressed with mu opioid receptors or whether agonists of one receptor can functionally alter the respiratory effects of the other.

Opioids, the most widely studied class of drugs to cause drug-induced respiratory depression, primarily depress respiration by eliciting changes in respiratory frequency; changes to tidal volume are observed inconsistently, and typically only at high doses or with high-potency synthetic agonists [28, 37–39]. Our results only partially recapitulate this pattern. CP55,940 suppressed tidal volume and frequency at the same dose in both males and females (0.1 mg/kg). THC suppressed frequency and tidal volume at the same dose in females (3 mg/kg), but at different doses in males (10 mg/kg and 1 mg/kg, respectively). CP55,940 generally produced respiratory suppressive effects at the same doses in both males and females, while THC generally produced effects at lower doses in females than in males. The topic of sex differences in drug overdose is an emerging one in the opioid literature. Population-level data from humans demonstrate that males are more likely to die of opioid overdose than females, though studies have demonstrated that females exhibit increased sensitivity to the analgesic effects of opioids [40]. To our knowledge sex differences in cannabinoid overdose are unexplored in current literature, though THC and its active metabolites have been shown to exhibit increased potency in female animals in various models, while conversely the metabolites of CP55,940 remain largely unexplored [41]. More work is necessary to determine whether the sex differences observed here may be attributed to differential metabolism of these drugs in males in females, differences in CYP induction, or differences in CB_1_ receptor density or efficacy in relevant brain regions. The mechanisms underlying sex differences observed here are unknown and require further investigation.

THC’s effects on respiratory activity are striking given the widespread nature of its use and the sparse nature of human overdoses. A floor effect was observed in THC’s effect on minute ventilation in mice of both sexes, as both the 10 mg/kg and 30 mg/kg dose reduced minute ventilation to similar levels. This floor, roughly 50 mL/minute in males and 45 mL/minute in females, may indicate that the lack of fatal THC overdoses observed in humans may be attributable not to a lack of drug effect, but to limitations on drug efficacy that limit respiratory effects to levels that do not constitute a clinical emergency. Further supporting this hypothesis is the observation that no dose of THC recapitulated the levels of respiratory suppression seen in CP55,940, in which near-complete elimination of respiratory activity was observed at the highest tested doses. To this extent it is noteworthy that we did not observe deaths from cannabinoids in any of our studies. The doses of THC tested here were near the limits of solubility in our vehicle; while we cannot rule out the possibility that THC may be systematically capable of achieving levels of respiratory suppression equivalent to those induced by CP55,940, it is practically unlikely to occur outside of an experimental setting.

The respiratory effects of CP55,940 and THC were mediated by CB_1_ receptors. The CNS-penetrant CB_1_ inverse agonist AM251 completely reversed the effects of both CP55,940 and THC in both sexes, indicating that observed effects were entirely mediated by activity at the CB_1_ receptor. The peripherally restricted CB_1_ antagonist AM6545 partially, but reliably, attenuated the respiratory effects of CP55,940, and partially attenuated some of the respiratory effects of THC. This indicates that observed effects are primarily mediated by CB_1_ receptors in the CNS, but that peripheral receptors are still contributing. Two notable exceptions to this pattern were the extension expiratory pause, which was prevented completely by AM6545 in animals treated with both THC and CP55,940, and inspiratory pause, which was completely prevented in animals dosed with CP55,940 and partially prevented in animals dosed with THC. These results suggest that peripheral CB_1_ receptors may play a role in maintaining the transition between inspiration and expiration. CB_1_ receptors are sparsely expressed in carotid body chemoreceptors, peripheral components of the respiratory system that modulate respiratory activity by sensing the partial pressure of oxygen and carbon dioxide in blood entering the brain through the carotid artery, though the precise functionality of these receptors is unknown [42, 43]. In the presence of AM6545, suppressive effects of THC on peak inspiratory and expiratory flow occurred earlier, suggesting that blocking peripheral CB_1_ receptors exacerbated the effects of THC on inspiratory and expiratory flow.

Notably, in our study respiratory effects were observed at or below doses commonly used for behavioral testing, and in many cases at doses that do not consistently capitulate hypothermia, catalepsy, analgesia, and locomotor suppression - cardinal behavioral signs of CB_1_ activation [44–47]. These behaviors – referred to as the cannabinoid tetrad [48] – are commonly measured by tasks that may be readily impacted by respiratory failure (e.g., open field, bar test, tail-flick). It is difficult to conceive that a mouse respirating at a small fraction of its normal rate would be capable of maintaining normal levels of locomotion. Reciprocally, catalepsy may reinforce the respiratory effects of CB_1_ agonists. Changes to tidal volume in opioid-induced respiratory depression are attributable to changes in chest muscle rigidity [49–51], and the rigid paralysis commonly observed after the application of cannabinoid agonists may contribute to the changes in tidal volume observed here. This hypothesis is reinforced by our observation that AM6545 partially reversed changes to tidal volume in animals injected with CP55,940.

Normal breathing in both humans and mice is characterized by a three-phase rhythm: inspiration, post-inspiration, and expiration [52–56]. After a brief post-expiratory pause, the cycle begins again; under ideal circumstances, continuing uninterrupted in perpetuity. Breathing is controlled by a network of highly interconnected chemoreceptors and microcircuits in the brainstem. Broadly, the preBötC generates inspiration, the retrotrapezoid nucleus/parafacial respiratory group orchestrates expiration, and the post-inspiratory complex orchestrates the transition between inspiratory and expiratory phases [24, 57–59]. The activity of these circuits is modulated by input from a variety of reciprocally connected nuclei, including the pontine Kölliker-Fuse and lateral parabrachial nuclei, which modulate inspiration, the Bötzinger Complex, a major source of inhibition that mainly contains expiratory neurons, and the Nucleus Tractus Solitarius, which modulates respiratory parameters in response to peripheral signals [60–68]. Of these regions, the preBötC has become a significant focus of research, as it contains a population of excitatory “pacemaker” neurons that serve as central pattern generators whose activity is necessary for rhythmic inspiration to occur [69]. The blockade of glutamatergic signaling in the preBötC causes a loss of rhythmic inspiration, leading to infrequent low-amplitude breathing and sustained interruptions to the respiratory cycle [70]. Furthermore, the region is critically implicated in the respiratory suppressive effects of exogenous opioids: application of the mu opioid antagonist naloxone into the preBötC prevents fentanyl-induced respiratory depression in a variety of animal models [71, 72], and deletion of mu opioid receptors in the preBötC attenuates respiratory rate depression by morphine [73, 74].

Here we show that CB_1_ mRNA is expressed in the preBötC, and that CB_1_ agonists suppress respiratory frequency. Furthermore, CB_1_ mRNA in the preBötC was frequently colocalized to neurons expressing VGLUT2. These results suggest that, similar to opioids, CB_1_-induced depression of respiratory rate may be mediated by CB_1_-expressing glutamatergic neurons in the preBötC. However, the preBötC contains both inspiratory and non-inspiratory glutamatergic neurons. It is unknown what proportion of CB_1_-expressing glutamatergic neurons observed in our experiments directly contribute to inspiration (e.g., express NK1R or DBX1, specific markers for inspiratory neurons). Additionally, it’s unlikely that CB_1_ expression in the preBötC drives the changes to tidal volume observed in our experiments. Consequently, further work is required to establish the precise distribution and contribution of CB_1_-expressing neurons in the preBötC, as well as expression of CB_1_ in other brain regions responsible for respiratory control, such as the Nucleus Tractus Solitarius and Kolliker-Fuse Nucleus.

## 5. Conclusions

Collectively, our findings demonstrate that both phytocannabinoid and synthetic CB_1_ agonists induce respiratory depression in a dose-dependent manner, impacting both respiratory and frequency and tidal volume to affect extreme reductions in ventilation. These effects were completely reversed by the brain-penetrant CB_1_ inverse agonist AM251, and partially reversed by the peripherally-restricted CB_1_ antagonist AM6545, indicating that they were caused by activity at CB_1_ receptors in both the CNS and periphery. Finally, we demonstrate that CB_1_ receptors are expressed in brain regions known to be critical to respiratory control.

## Acknowledgements

This work was supported by National Institutes of Health National Institute on Drug Abuse (NIDA) grant DA09159 (AGH), Frontier Program KKP129961 (IK), National Institutes of Health grant R01DA044925 (IK), and T32 NIDA Predoctoral Training Grant DA024628 (JW).

---

**Supplemental 1:**
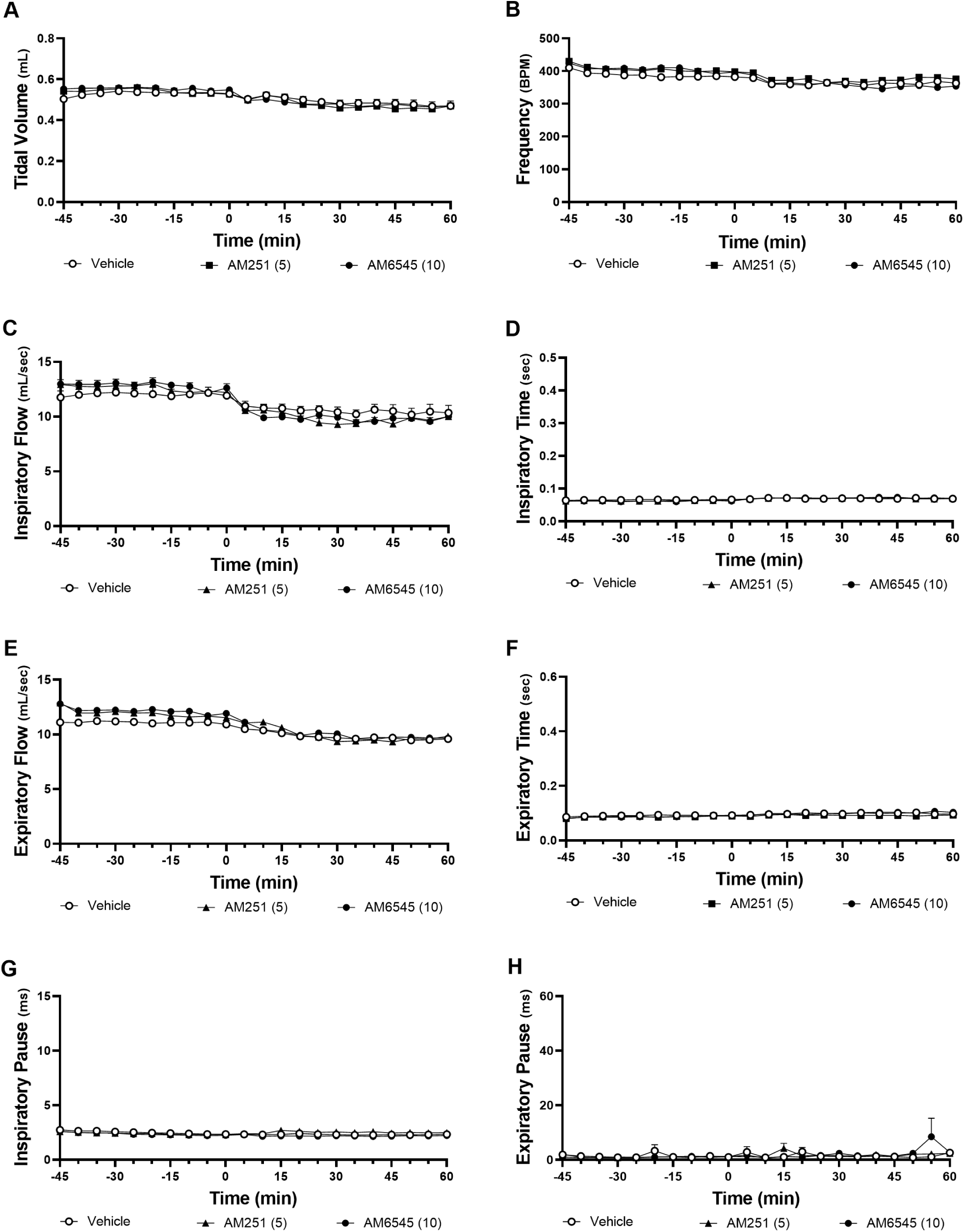
Antagonists Alone non-Minute Ventilation

